# An analysis of avian vocal performance at the note and song levels

**DOI:** 10.1101/664896

**Authors:** David M. Logue, Jacob A. Sheppard, Bailey Walton, Benjamin E. Brinkman, Orlando J. Medina

## Abstract

Sexual displays that require extreme feats of physiological performance have the potential to reliably indicate the signaller’s skill. The hypothesis that the structure of bird song is physiologically constrained remains controversial. We tested for evidence of performance constraints in Adelaide’s warblers (*Setophaga adelaidae*) songs. At the note level, we identified three trade-offs with well-defined limits. At the song level, we identified two trade-offs, but their limits were less well-defined than the note-level limits. Trade-offs at both levels suggest that song structure is constrained by limits to the speed of both frequency modulation (while vocalizing and between notes) and respiration. Individual males experience the same trade-offs that characterize the population, but the intensity of those trade-offs varies among individuals. Performance metrics derived from the observed limits to performance varied moderately among individuals and strongly among song types. Note-level performance metrics were positively skewed, as predicted by the hypothesis that this population has experienced positive selection for constrained performance. We conclude that physiological limits on frequency modulation and respiration constrain song structure in male Adelaide’s warblers. Further work is needed to determine whether receivers respond to natural levels of variation in performance, and whether performance correlates with singer quality.

## Introduction

Animal displays are dynamic signals that often function as sexual advertisements. Many sexual displays challenge the signaller’s motor abilities (Byers et al. 2010). For example, male hummingbird courtship displays are pinnacles of length-specific velocity and acceleration (Larimer and Dudley 1995; Clark 2009), male pronghorns’ (*Antilocapra americana*) circle chase displays require extreme angular acceleration (Byers 1997), and the success of male fiddler crabs’ (*Uca perplexa*) displays depends on the speed and height of their waving claws (Murai and Backwell 2006). In these examples and others, it appears that female choice for high performance has promoted the evolution of displays that reliably showcase males’ abilities to perform near their species’ physiological limits (Smith and Harper 2003; Byers et al. 2010; Bradbury and Vehrencamp 2011). Bird song is another sexual display that may fit that description (Nowicki et al. 1992; Podos 1997; Podos et al. 2009).

### Performance constraints in bird song

Song performance has been proposed to serve as a reliable signal of sender condition that is salient to both male and female receivers (reviewed in Podos 2017). To sing, a bird must rapidly coordinate the output of its brain, syrinx, bill, and respiratory system. Motor constraints on any of these systems, or on the ability to coordinate them, could generate reliable correlations between motor abilities and song structure, and signal receivers could learn about a signaller’s quality by listening to his or her song. A history of female choice for (or male deference to) males that sing with high performance could explain the widespread evolution of songs with rapid changes in fundamental frequency and rapid sequences of notes.

Several physiology studies indicate that motor constraints limit song structure (Hoese et al. 2000; Podos and Nowicki 2004; Plummer and Goller 2008). Most studies of motor performance in bird song, however, do not directly measure physiological limits during song production (Cardoso 2017). Rather, they test the prediction that physiological constraints induce trade-offs between acoustic traits. Podos (1997) performed the first such analysis when he described a trade-off between the trill rate and frequency bandwidth in sparrow (Emberizidae) songs. Sparrow songs with wide frequency bandwidth do not have high trill rates, and those with high trill rates do not have wide frequency bandwidths (Fig. 5 in Podos 1997). This pattern can be at least partially explained by limits on the speed of voiced frequency modulation (FM) and unvoiced FM (frequency jumps between notes). Since Podos’s pioneering work, several other studies have identified trill rate vs. frequency bandwidth trade-offs in other species (reviewed in Wilson et al. 2014; Podos 2017).

**Figure 5:**
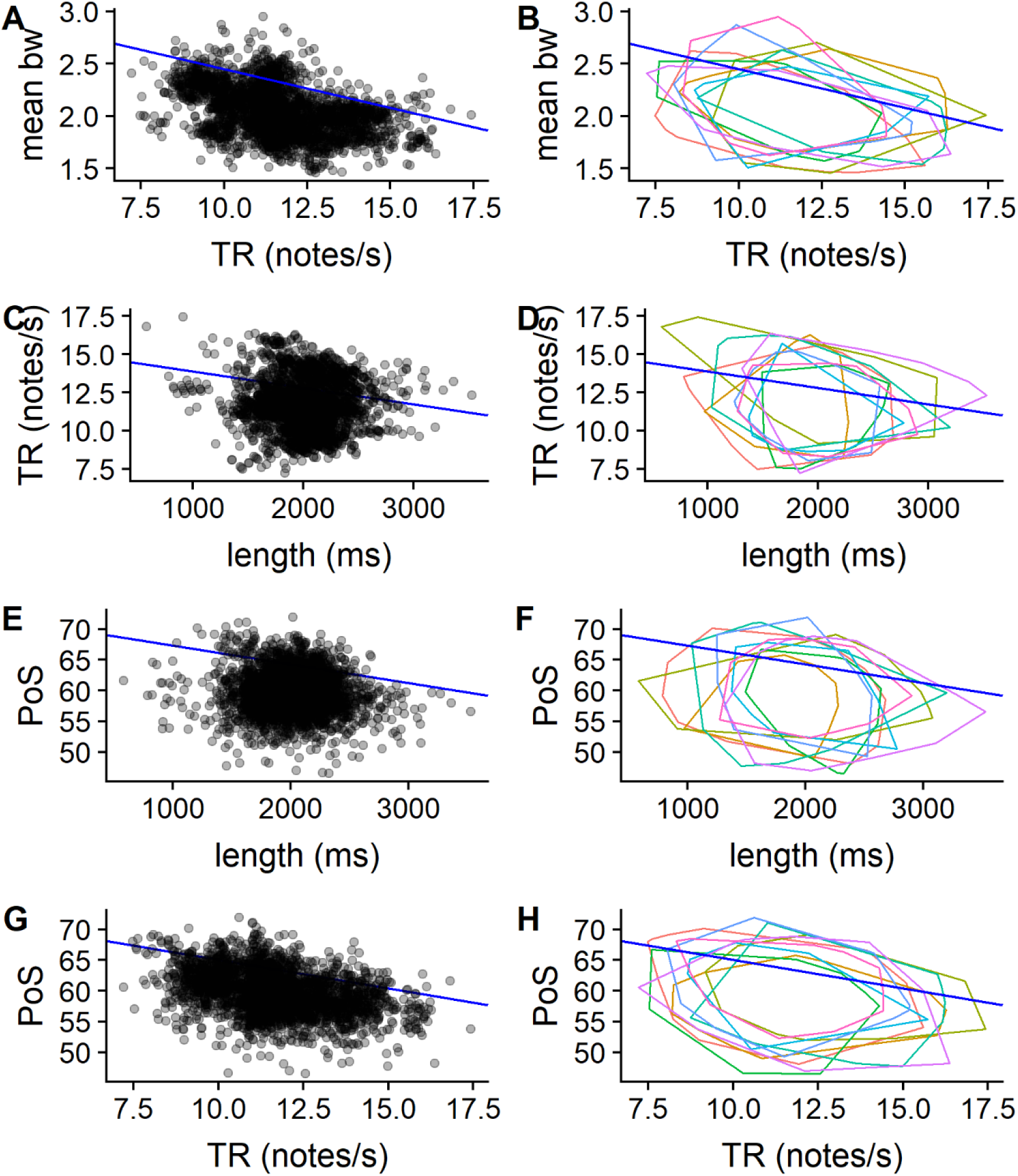
Plots comparing acoustic properties of Adelaide’s warbler songs. Data points in scatterplots (A, C, E, G) are semi-transparent to facilitate interpretation. Each colored polygon in the polygon plots (B, D, F, H) represents the limits of an individual males’ song sample. Royal blue lines represent 90^th^ quantile regression lines.

Bird song researchers have identified several other acoustic trade-offs that suggest performance constraints since Podos’s studies of trill rate vs. frequency bandwidth trade-offs. A few have started to study performance constraints at the level of the note. Relative to song-level analyses, note-level analyses can be expected to provide clearer evidence of trade-offs if there is less unmeasured structural variation among notes than there is among songs. For a given number of recordings, note-level analysis also produces more data points than song-level analysis, resulting in a more thorough description of acoustic space. Finally, note-level analyses permit tests of hypotheses about specific physiological constraints, because they allow researchers to isolate song elements related that might be subject to that constraint.

A recent note-level study showed that the difference in fundamental frequency between the end of one note and the beginning of the next note trades off against the length of the silent gap between notes (Geberzahn and Aubin 2014a). Larger frequency jumps between syllables require longer silent gaps. This trade-off suggests a limit to the speed of unvoiced FM, one of the constraints that may contribute to the widely-observed trade-off between trill rate and frequency bandwidth.

A second line of note-level research showed that longer notes are associated with longer silent gaps before the subsequent note (Mota and Cardoso 2001; Cardoso et al. 2007). This trade-off seems to be caused by a respiratory constraint. When singing rapidly, songbirds take unvoiced mini-breaths between notes (Hartley and Suthers 1989). Long notes use more air than short notes, so they require longer subsequent mini-breaths (Suthers and Zollinger 2004). Extremely short notes are produced by a mechanism called ‘pulsatile expiration,’ which does not require mini-breaths (Hartley and Suthers 1989).

### Measuring performance in bird song

Trade-offs in acoustic traits can be used to quantify song performance. Cardoso (2017, p. e29) defines song performance as ‘the degree of challenge to the motor system, the respiratory system, or other physiological processes involved in singing’ (see also Byers et al. 2010). Song performance metrics based on acoustic trade-offs estimate performance based on the acoustic distance to an observed acoustic performance limit (Podos 2001). Vocalizations that are close to or beyond the limit are high performance, whereas those that are far from the limit are low performance (Fig. 1). Performance measurements derived in this way are called ‘deviations’ because they measure the orthogonal deviation from the performance limit (Podos 2001).

**Figure 1:**
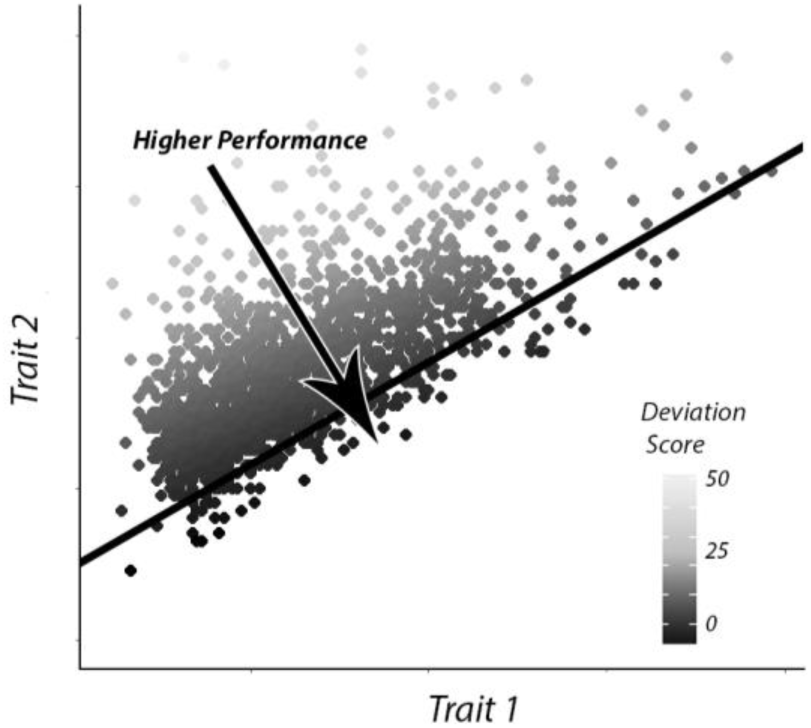
Scatterplot depicting a hypothetical performance constraint between two traits. The 10^th^ quantile regression line is shown in black. For comparisons in which the trait on the x-axis scales positively with performance and the trait on the y-axis scales negatively with performance, any performance limit would follow the lower edge of the distribution and slope upward, and performance would increase as one moves down and to the right (this figure, Fig. 4). For comparisons in which both the x- and y-axes scale positively with performance, performance limits follow the upper edge of the distribution and slope downward, and performance increases as one moves up and to the right (e.g., Fig. 5).

### Critiques of bird song performance research

Recently, research on bird song performance has come under criticism (Kroodsma 2017b, 2017a). Kroodsma argues that performance is unlikely to be a reliable signal of singer quality because it varies a great deal among song types, but little or not-at-all among individuals. One way to test this hypothesis is to measure the repeatability (intraclass correlation coefficient, ICC) of putative performance metrics for individuals and for song types (Cardoso et al. 2009). We want to know whether a receiver’s ability to compare individual singers’ performance is limited by the ICC for individuals, so our ICC calculation should reflect the receiver’s assessment strategy. The ICC for individuals should be based on the average performance of each song type from each singer (as in Cardoso et al. 2009) if receivers first assess each song type separately, and then average the performance of all of the song types in a singer’s repertoire to estimate his quality. However, if receivers assess singers’ performance based on all of the songs they hear, without first averaging within song type, the ICC should be based on all sampled songs. We did not know how (or whether) receivers assess singers’ performance, so we applied both methods.

Kroodsma (2017b) also argues that the bounded scatterplots that others have interpreted as evidence of performance constraints (e.g., Figs. 1, 4, & 5) do not represent performance limits, but are a consequence of the cultural transmission of song structure. According to this ‘constrained learning’ hypothesis, it is physically possible for birds to sing beyond the observed limits, but they do not because they have not learned songs beyond these limits. In any specific case, the performance constraint hypothesis and constrained learning hypothesis are mutually exclusive because they attempt to explain the same thing (bounded scatterplots). We used two predictions to distinguish between these competing hypotheses. First, we predicted that performance constraints would result in scatterplots with boundaries that slope in the expected direction given plausible motor constraints. Constrained learning may or may not result in sloped boundaries, and the direction of the slope should be random with respect to plausible motor constraints. Second, if a performance limit constrains the structure of vocalizations and singers have evolved under positive selection for singing performance, vocalizations should cluster near the limit, and deviation scores should skew away from the limit. The constrained learning hypothesis generates a different prediction: Selection for species-specific song structure should cause vocalizations to cluster in the middle of the distribution, where they are least likely to be mistaken for heterospecific song. This pattern would produce a symmetrical deviation distribution (skew ≈ 0) with diffuse (low-density) edges.

### The present study

We analysed a sample of male Adelaide’s warbler (*Setophaga adelaidae*) songs for evidence of performance constraints. We tested for three trade-offs at the note level and four trade-offs at the whole song level, while controlling for variation attributable to individuals. We then analysed the deviation scores. We tested whether receivers might be able to distinguish high-performance singers from low-performance singers by estimating note-level and song-level ICC’s for individuals. We also estimated ICC’s for song types, to better understand the relationship between song type repertoires and vocal performance, and to permit comparisons with other species. Finally, we tested opposing predictions of the constrained learning and constrained performance hypotheses by measuring the skewness of the deviation distributions. Although several previous studies have examined acoustic trade-offs in bird songs, this study is distinguished by its comprehensiveness, sample size at the note level, and novel analytic approach.

## Methods

### Study species

Adelaide’s warbler is a socially monogamous insectivore endemic to Puerto Rico and Vieques. Mated pairs maintain all-purpose territories throughout the year (Toms 2010). Male songs are frequency modulated trills (Fig. 2). Individual males sing repertoires of song types (avg. ≈ 23 songs), many of which are shared with other males (Staicer 1991; Kaluthota et al. 2019). Each male’s repertoire comprises two categories, A and B, which are characterized by distinct times of delivery, song rates, and song switching frequencies (Staicer 1991; Kaluthota et al. 2019). The individual notes comprising songs are structurally simple, with almost all of the energy concentrated in the fundamental frequency. There exists considerable among-note variation in length, frequency, and frequency modulation (Fig. 2).

**Figure 2:**
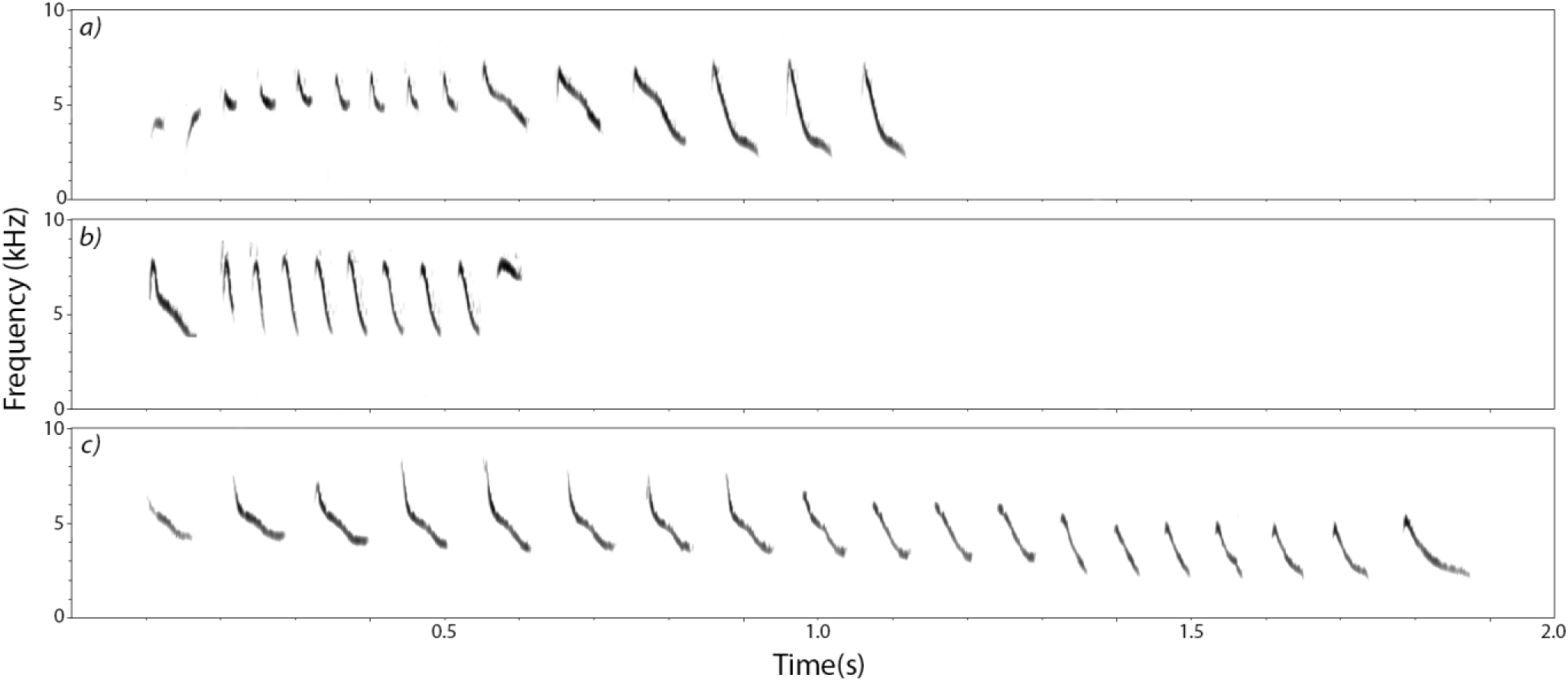
Sound spectrograms of three Adelaide’s warbler song types.

### Ethics

This research was approved by the Institutional Animal Care and Use Committee at the University of Puerto Rico, Mayagüez (17 September, 2010). It adheres to the ASAB/ABS Guidelines for the use of animals in research. Birds were captured under D.M.L.’s federal bird banding permit (no. 23696). The U.S. Fish and Wildlife Service granted permission to work at the Cabo Rojo Wildlife Refuge (permit 2012-01).

### Recording and scoring

We recorded nine mated male Adelaide’s Warblers at the Cabo Rojo National Wildlife Refuge (US Fish and Wildlife Service) during the breeding season between March and June, 2012. Each male was recorded on four days, from 45 minutes before sunrise until 2 hours and 45 minutes after sunrise. Consecutive recordings of a given male were separated by at least four days, except on two occasions when recordings were made on consecutive days because of logistical constraints. Recordings were made with Marantz PMD 661 digital recorders and Sennheisser ME67 shotgun microphones (file format = wav, sampling rate = 44.1KHz, bit depth = 16 bits). This is the same set of recordings used in a previous study of short-term variation in song performance (Schraft et al. 2017), a methods paper on song sequences (Hedley et al. 2018), and an analysis of singing modes (Kaluthota et al. 2019).

We visualized recordings as sound spectrograms in Syrinx PC v.2.6 (J. Burt, Seattle, WA; Settings: Blackman window, transform size = 1024 points). Each song recording from a focal male was assessed for recording quality. One person (D.M.L.) assigned song recordings to song types. The inter-rater reliability of song type scoring of these recordings was estimated to be 100% within an individual bird, and 87% among individuals (Kaluthota et al. 2019). We only used high-quality recordings (high signal-to-noise ratio, minimal overlap with other sounds) for song measurements. High quality song recordings were analysed in Luscinia v2.14 (max. freq. = 10kHz, frame length = 5ms, time step = 1ms, dynamic range = 35 dB, dynamic equalization = 100ms, de-reverberation = 100%, de-reverberation range = 100ms, high pass threshold = 1.0kHz, noise removal = 10dB; Lachlan 2007). We loosely outlined the trace of each note with a stylus on a touchscreen monitor, and Luscinia’s algorithms identified the signal and rejected background noise. Acoustic parameters for all notes were output to a spreadsheet. Luscinia offers several frequency metrics. We chose peak frequency because visual inspection of spectrograms showed that it tracked the fundamental frequency better than the fundamental frequency metric.

### Analysis: trade-offs

The note-level analysis omitted the one or two low-amplitude notes that began some songs and the final note of all songs. We omitted final notes because it was impossible to define the length of the silent gap following the last note. For the note-level analyses, we analysed the frequency bandwidths and durations of both notes and silent gaps (Fig. 3). Following Cardoso (2013), our measures of frequency bandwidth (BW) comprise the ratio of the maximum frequency to the minimum frequency. Gap length and gap BW are taken from the silent gap after the focal note. Gap BW is based on the end of the focal note and the beginning of the subsequent note. We tested three comparisons at the note level that might reveal trade-offs indicative of performance constraints: note length vs. gap length (respiratory), note BW vs. note length (voiced FM), and gap BW vs. gap length (unvoiced FM).

**Figure 3.**
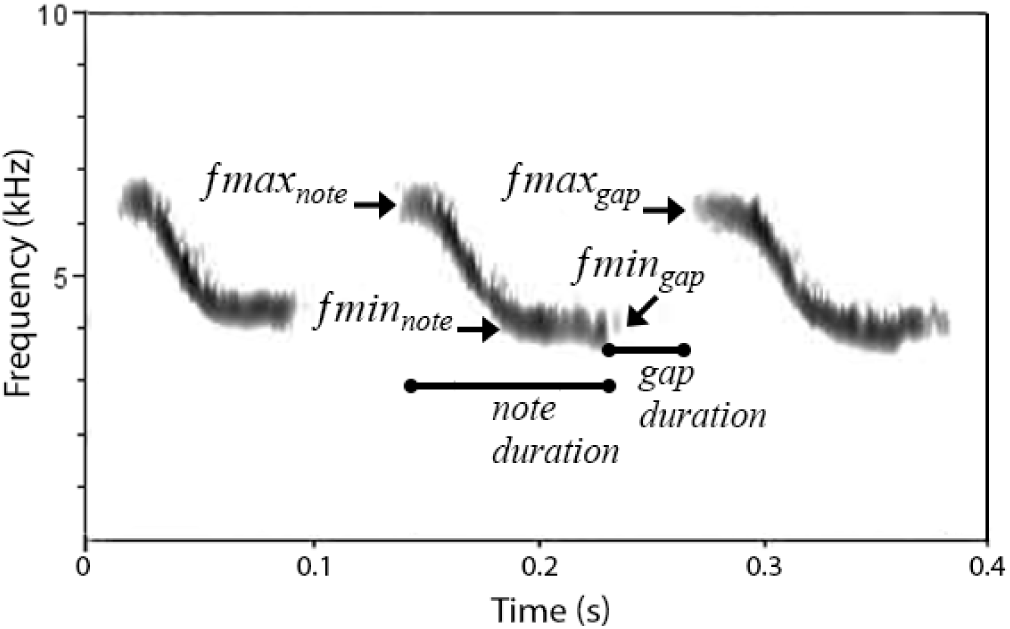
Six note-level measurements. The focal note is in the middle. We used *Ƒmax_note_* and *Ƒmin_note_* to determine the note frequency bandwidth, and *Ƒmax_gap_* and *Ƒmin_gap_* to determine the gap frequency bandwidth.

We chose four parameters for the song-level analyses: trill rate, average frequency bandwidth, percent of sound, and duration. Trill rate (TR) was calculated as the number of notes in the song minus one, divided by the time from the beginning of the first note to the beginning of the last note. We excluded the final note from this calculation because TRs based on the full song necessarily omit the ‘gap’ after the last note, biasing estimates upward for songs with fewer notes. Adelaide’s warbler songs are frequency modulated trills (Fig. 2), so the total frequency bandwidth of a song is only weakly related to the amount of FM in the song. We therefore calculated a song’s BW as the average BW of the notes in the song. Percent of sound (PoS) is the percent of the song that is voiced. It was calculated as the sum of note lengths, divided by the total length of the song, multiplied by 100. Lastly, we measured song length because many kinds of performance increase in difficulty with increasing duration (Byers et al. 2010). We tested four comparisons that might reveal trade-offs at the song level: TR vs. mean BW (FM, respiratory), length vs. TR (respiratory endurance), length vs. PoS (respiratory endurance), and TR vs. PoS (respiratory).

Wilson et al. (2014) recommend quantile regression to test for acoustic trade-offs. Quantile regression produces a linear function to estimate a defined quantile Y over a range of X (Cade and Noon 2003). We conducted mixed quantile regression analyses to characterize potential trade-offs while accounting for the non-independence of acoustic units from the same individual (Geraci 2014). To the best of our knowledge, this is the first study to use mixed quantile regression to test for acoustic trade-offs. To conform to recommended best practices, we treated both intercepts and slopes as random variables (Barr et al. 2013). We predicted positive lower boundaries for all note-level analyses, because higher values of acoustic variable X were predicted to constrain the minimum values of variable Y (as in Geberzahn and Aubin 2014a). In contrast, we predicted negative upper boundaries for all song-level analyses, because higher values of variable X were predicted to constrain the maximum values of variable Y as in (as in Podos 1997). For the note-level dataset, we tested lower boundaries with 10^th^ quantile regressions (tau = 0.10), following advice in Wilson et al. (2014). Similarly, we ran 90^th^ quantile regressions (tau = 0.90) to estimates upper boundaries in the song-level dataset (Wilson et al. 2014).

Population-level performance limits could arise from a pooled analysis of individuals that do not themselves exhibit trade-offs. For example, some individuals might sing high-trill-rate, low-bandwidth songs while others sing low-trill-rate, high-bandwidth songs, resulting in a sloping limit to the population’s distribution when individuals’ data are pooled. Alternatively, different individuals may be subject to similar trade-offs, which combine to produce a population-wide trade-off. We therefore examined data from individual birds for evidence of trade-offs. We also asked whether different individuals experience trade-offs differently by generating a reduced model without random slopes and comparing its Akaike Information Criterion (AIC) value against that of the full model. Differences in AIC values (ΔAIC) greater than 20 were interpreted as evidence that the model with the higher AIC lacked support (Burnham et al. 2010).

In addition to the hypothesis tests described above, we offer several visual representations of our data. We graphed the whole distributions with semi-transparent points and average quantile regression lines. To show individual variation near the boundaries, we also present zoomed-in views of the boundary regions with separate colours for each individual and polygons that mark the limit of each individual’s distribution. Polygons were generated with the *geom_encircle* command in the *ggalt* package (Rudis et al. 2017), with settings s_shape =1 (no added curvature) and expand = 0 (polygon edges intersect extreme points). Finally, we present separate distributions for each individual in the electronic supplementary material.

### Analysis: Performance metrics

Deviation scores were calculated as the orthogonal distance from the quantile regression lines, such that higher (more positive) values indicate lower putative performance (Podos 2001). We estimated intra-class correlations (ICCs) to test how repeatable individuals were with respect to note-level deviation scores averaged over songs and song-level deviation scores. We conducted ICCs for individuals using both the entire sample and the average scores for each song type within individual (see Introduction). We also tested the repeatability of song types for note-level deviation scores averaged over songs and song-level deviation scores. We used log-likelihood tests to generate p-values for the ICCs.

We generated Pearson’s correlation matrices of deviation scores for notes and songs. The song-level correlation matrix included song-level deviations, note-level deviations averaged over songs, and the performance metric Frequency Excursion (FEX). Frequency excursion attempts to estimate the speed with which the vocal apparatus adjusts frequency by measuring the rate of change in the fundamental frequency, including during silent gaps (Podos et al. 2016; Schraft et al. 2017). We calculated the skewness of the deviation scores to test competing predictions of the constrained performance and constrained learning hypotheses.

All statistics were conducted in R Studio (Team 2015). Mixed quantile regressions relied on the *lqmm* package (Geraci 2014). Intra-class correlations were assessed with the *rptR* package (Stoffel et al. 2017). Data visualizations relied on the package *ggplot2* (Wickham and Chang 2008). Data and R code are available at (*DRYAD link when available*).

## Results

### Note-level: Descriptive statistics

Subjects contributed an average of 320 ± 185 (average ± SD) songs, belonging to 20.8 ± 3.7 song types and comprising 7622 ± 4568 notes, to our analyses (Table 1). On average, notes were 50.8 ± 23.9 ms long and silent gaps were 35.7 ± 11.5 ms long. Our measure of note-level frequency bandwidth, the ratio of maximum peak frequency to minimum peak frequency, averaged 2.0 ± 0.57 for notes, and 1.93 ± 0.57 for silent gaps.

**Table 1.**
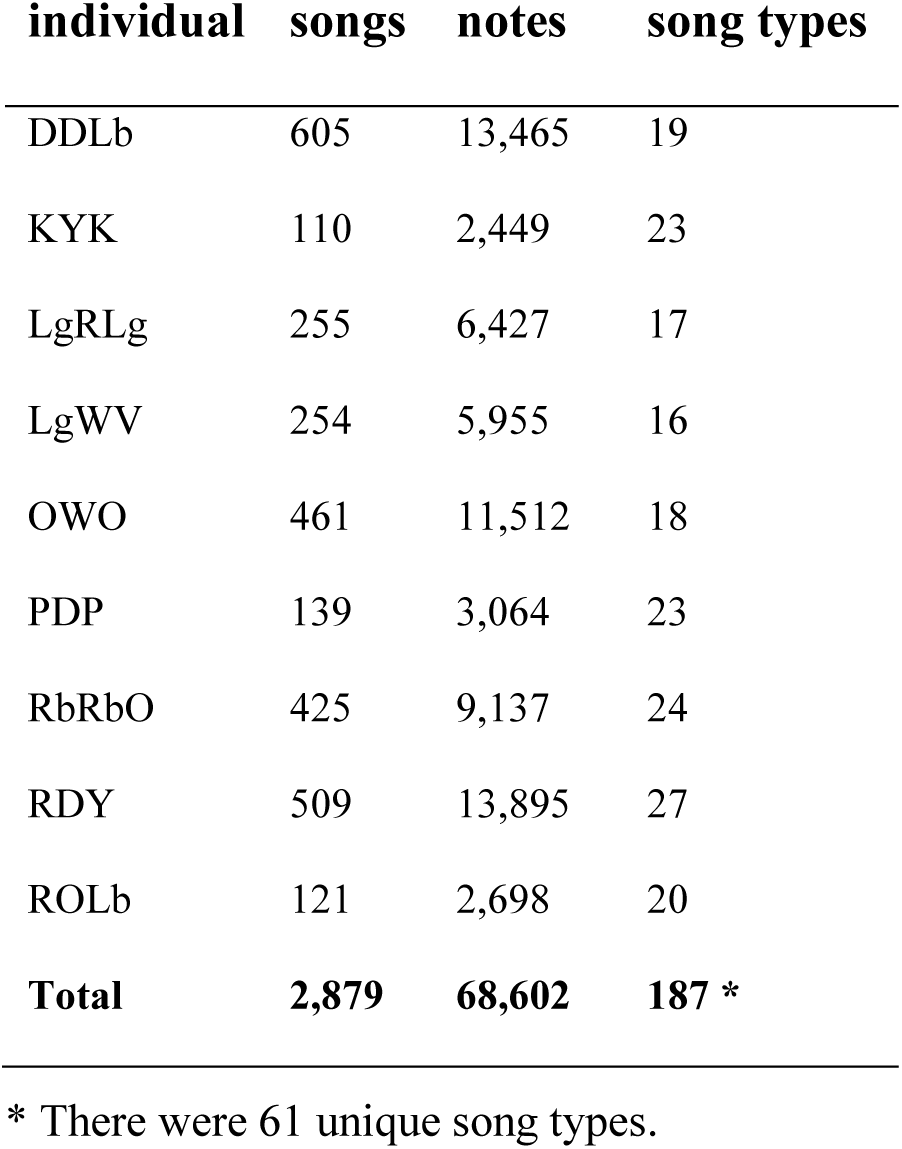
Sample sizes, separated by individual.

### Note level: Trade-offs

Mixed quantile regression analyses for note length vs. gap length (intercept = 12.87, slope = 0.24, p_slope_ < 0.0001) and note BW vs. note length (intercept = 18.45, slope = 8.46, p_slope_ < 0.0001) were significant and positive, and all individuals’ slope estimates were positive (Table ESM-1). The analysis of gap BW vs. gap length produced a positive slope estimate (intercept = 23.02, slope = 3.19, p_slope_ = 0.03), but the p-value was marginally significant at the α = 0.05 level, and eight out of nine individuals’ slope estimates were positive. Individuals’ slopes varied significantly in the note length vs. gap length (ΔAIC = 981) and gap BW vs. gap length (ΔAIC = 2724) comparisons, but not in the note BW vs. note length comparison (ΔAIC = −178).

Pooled scatterplots revealed sharply demarcated, approximately linear lower boundaries, except for a prominent bulge in the lower left part of the boundary on the note length vs. gap length plot (Fig. 4a-c). Average quantile regression lines approximated the slopes of boundaries of the distributions, but their fit was imperfect (Fig. 4). The fit of the average quantile regression line was particularly poor for the gap BW vs. gap length distribution (Fig. 4c). The polygons and individual-level scatter plots showed positively sloping lower boundaries for all individuals, and inter-individual variation in the slopes of the boundaries (Figs. 4d-f; ESM-1-3). The bulge in the lower left portion of the note length vs. gap length comparison is apparent in most of the individual-level plots (Fig. ESM-1).

**Figure 4:**
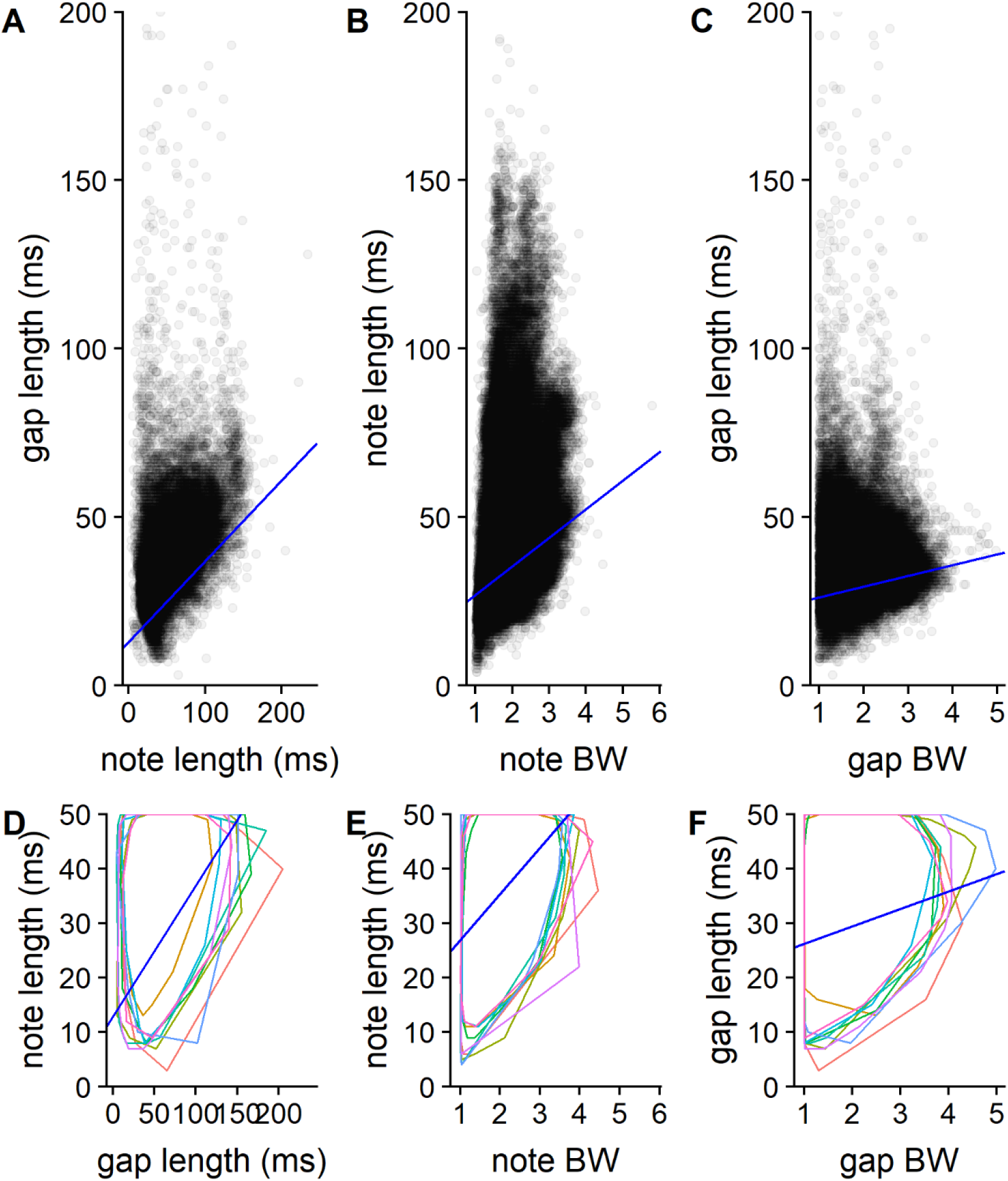
Plots comparing acoustic properties of Adelaide’s warbler notes. Data points in scatterplots (A-C) are semi-transparent to show density. Polygon plots (D-F) focus on the bottom of each distribution. Each colored polygon represents the limits of an individual males’ note sample. Royal blue lines represent 10^th^ quantile regression lines. Scales differ between the scatterplots and the polygon plots.

### Note-level: Deviations

When we used the entire sample of songs, all three note-level deviations averaged over songs were significantly repeatable by individual, with moderate repeatability estimates (Table 2). Using the average deviation scores for each song type from each individual resulted in lower repeatability estimates, and rendered the note BW vs. note length deviation ICC non-significant at the α= 0.05 level. All three deviation scores were significantly repeatable among song types, with moderate to high repeatability estimates.

**Table 2.**
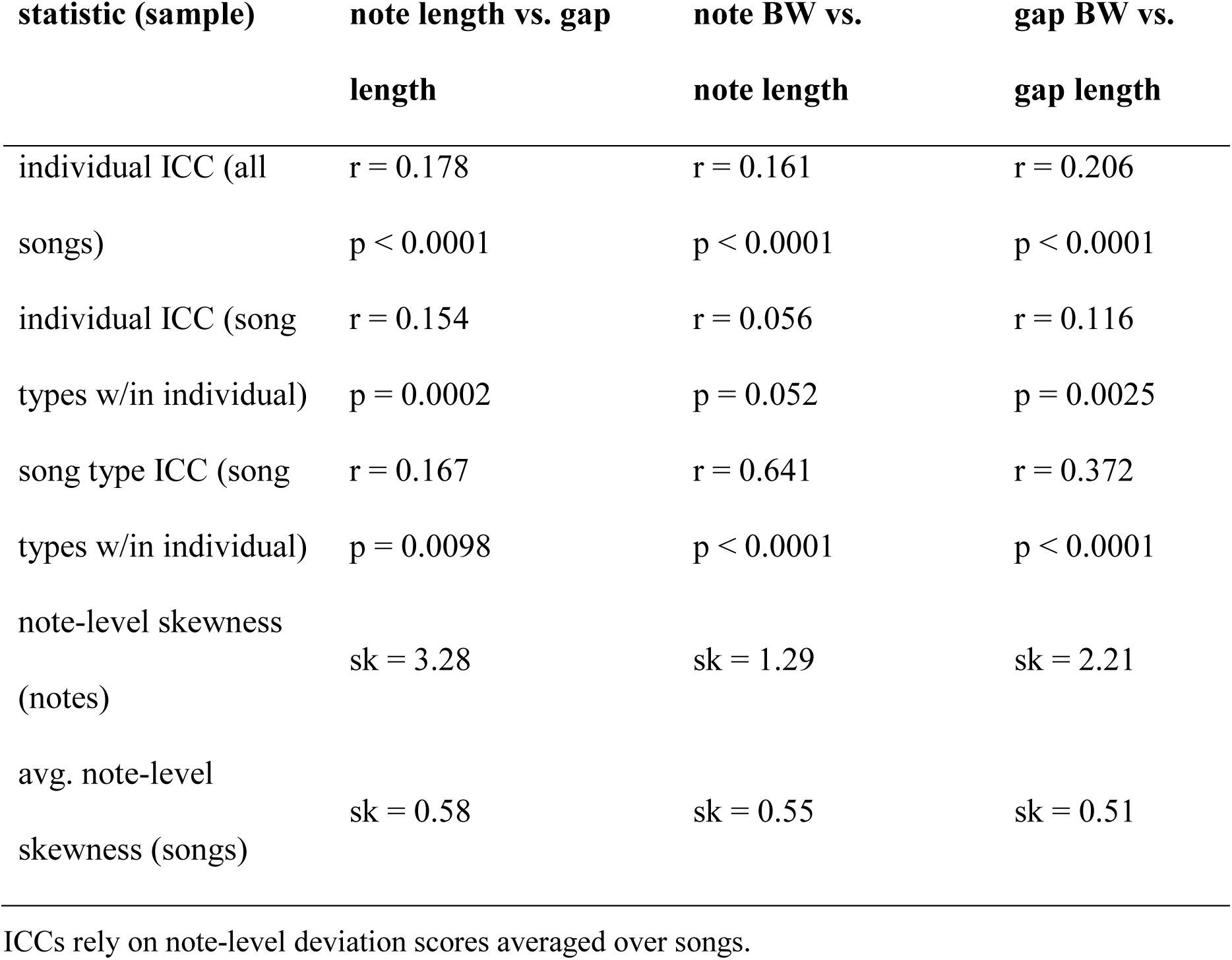
Intra-class correlation and skewness values for note-level deviation scores.

| statistic (sample) | note length vs. gap<br>length | note BW vs.<br>note length | gap BW vs.<br>gap length |
| --- | --- | --- | --- |
| individual ICC (all<br>songs) | $r = 0.178$<br>$p < 0.0001$ | $r = 0.161$<br>$p < 0.0001$ | $r = 0.206$<br>$p < 0.0001$ |
| individual ICC (song<br>types w/in individual) | $r = 0.154$<br>$p = 0.0002$ | $r = 0.056$<br>$p = 0.052$ | $r = 0.116$<br>$p = 0.0025$ |
| song type ICC (song<br>types w/in individual) | $r = 0.167$<br>$p = 0.0098$ | $r = 0.641$<br>$p < 0.0001$ | $r = 0.372$<br>$p < 0.0001$ |
| note-level skewness<br>(notes) | $sk = 3.28$ | $sk = 1.29$ | $sk = 2.21$ |
| avg. note-level<br>skewness (songs) | $sk = 0.58$ | $sk = 0.55$ | $sk = 0.51$ |

ICCs rely on note-level deviation scores averaged over songs.

Deviation scores from the gap BW vs. gap length and note length vs. gap length comparisons were highly positively correlated (r = 0.874). There was a moderate correlation between gap BW vs. gap length and note BW vs. note length scores (r = 0.574), and a weak correlation between note BW vs. note length and note length vs. gap length deviations (r = 0.112).

The distributions of deviation scores were all positively (right) skewed (Table 2). Note-level deviation scores were strongly skewed, while note-level deviations averaged over songs produced moderately skewed distributions. We graphed deviation against rank to identify statistical outliers that fell below the regression lines (n = 72). Most of these outliers were attributable to either reverberation or background noise that was accidentally treated as signal when the songs were outlined in Luscinia, resulting in two notes being merged into one. We excluded these outliers from graphs, but we did not exclude them from statistical analyses because they comprise a small proportion of the overall dataset (∼0.1%), and pruning only those scoring errors that were outliers would bias the dataset.

### Song level: Descriptive statistics

TRs averaged 11.64 ± 1.58 notes / second. Average mean frequency bandwidths were 2.05 ± 0.23. The average PoS was 59.80 ± 3.52 %. Songs were 2025.56 ± 299.11 ms long, with an FEX score of 66.39 ± 11.45, on average.

### Song level: Trade-offs

Quantile regression analyses indicated the presence of statistically significant, negatively sloping upper boundaries for all four song-level comparisons, although the TR vs. PoS slope was only marginally significant (TR vs. mean BW: intercept = 3.19, slope = −0.074, p_slope_ < 0.0001; length vs. TR: intercept = 14.95, slope = −0.0011, p_slope_ = 0.0071; length vs. PoS: intercept = 70.38, slope = −0.003, p_slope_ = 0.0001; TR vs. PoS: intercept = 74.21, slope = −0.92, p_slope_ = 0.0356). All of the individual slope estimates were negative except for one individual’s length vs. TR estimate (Table ESM-2). Individuals’ slopes varied significantly in the TR vs. mean BW (ΔAIC = 3529), length vs. TR (ΔAIC = 4648), and TR vs. PoS (ΔAIC = 8473) comparisons, but not in the length vs. PoS comparison (ΔAIC = −23,945).

The pooled scatterplots suggest overall negative trends and loosely-defined negatively sloping upper limits for the TR vs. mean BW and TR vs. PoS comparisons (Fig. 5a, g). The average 90^th^ quantile regression lines appeared to fit the upper boundaries of the TR vs. mean BW and TR vs. PoS distributions well (Fig. 5a, g). Polygons and individual-level scatterplots from most individuals showed evidence of negative relationships for the TR vs. mean BW and TR vs. PoS comparisons (Figs. 5b, h, ESM-4, ESM-7). In contrast, the length vs. TR and length vs. PoS pooled scatterplots, polygons, and individual-level scatterplots did not provide evidence of negative relationships (Fig. 5c-f, ESM-5, ESM-6).

### Song level: Deviations

All four song-level deviation scores and FEX were moderately repeatable among individuals when all songs were considered (Table 3). When we used the average of each song type within individual to calculate ICCs, all repeatability estimates were lower except for those derived from the TR vs. PoS deviations, and the TR vs. mean BW repeatability was not statistically distinguishable from zero. The song-level metrics of performance were moderately to highly repeatable among song types.

**Table 3.** Results of intra-class correlation analysis and skewness analysis on song-level deviation scores and FEX.

| statistic (sample) | TR vs.<br>mean<br>BW | length vs.<br>TR | length vs.<br>PoS | TR vs.<br>PoS | FEX |
| --- | --- | --- | --- | --- | --- |
| individual ICC (all songs) | r = 0.161<br>p < 0.0001 | r = 0.158<br>p < 0.0001 | r = 0.156<br>p < 0.0001 | r = 0.144<br>p < 0.0001 | r = 0.140<br>p < 0.0001 |
| individual ICC (song types w/in individual) | r = 0.009<br>p = 0.455 | r = 0.079<br>p = 0.011 | r = 0.126<br>p = 0.0006 | r = 0.148<br>p = 0.0002 | r = 0.068<br>p = 0.040 |
| song type ICC (song types w/in individual) | r = 0.556<br>p < 0.0001 | r = 0.604<br>p < 0.0001 | r = 0.280<br>p < 0.0001 | r = 0.117<br>p = 0.0254 | r = 0.534<br>p < 0.0001 |
| skewness (all songs) | -0.092 | -0.264 | 0.162 | 0.138 | -0.070 |

FEX, song-level deviations, and note-level deviations averaged over songs were intercorrelated (Table 4). We found strong negative relationships between FEX and each of the following deviation scores: TR vs. mean BW, mean note BW vs. note length and mean gap BW vs. gap length. Rapid FM and high TRs are indicated by high FEX values and low deviations, so these negative correlations may imply redundancy. By contrast, FEX was only weakly correlated with the TR vs. PoS and mean note length vs. gap length deviations. The song level metric TR vs. PoS deviation correlated strongly with the mean note-level deviations from the note length vs. gap length and gap BW vs. gap length deviations. The mean gap BW vs. gap length deviations were also highly correlated with the mean note length vs. gap length deviations and the mean note BW vs. note length deviations, but those two were not strongly correlated with each other.

**Table 4.** Pearson’s correlation matrix for song-level deviation scores, frequency excursion, and note-level deviation scores averaged over songs.

|  | TR vs.<br>mean<br>BW | length<br>vs. TR | length<br>vs. PoS | TR vs.<br>PoS | mean note<br>length vs.<br>gap length | mean note<br>BW vs. note<br>length | mean gap<br>BW vs.<br>gap length |
| --- | --- | --- | --- | --- | --- | --- | --- |
| **FEX** | -<br>0.707 | -0.310 | 0.072 | -<br>0.063 | -0.229 | -0.420 | -0.423 |
| **TR vs. mean BW** |  | 0.108 | 0.108 | 0.156 | 0.187 | 0.223 | 0.310 |
| **length vs. TR** |  |  | -<br>0.343 | 0.017 | 0.416 | 0.884 | 0.728 |
| **length vs. PoS** |  |  |  | 0.873 | 0.636 | -0.638 | 0.282 |
| **TR vs. PoS** |  |  |  |  | 0.886 | -0.293 | 0.640 |
| **mean note length**<br>**vs. gap length** |  |  |  |  |  | 0.090 | 0.900 |
| **mean note BW**<br>**vs. note length** |  |  |  |  |  |  | 0.508 |

## Discussion

Sexual displays can reliably indicate the signaller’s skill when physiology constrains display structure. We found evidence that the structure of a bird’s song is constrained by the speed with which singers can modulate frequency and replenish expired air. Some males consistently performed closer to the estimated population-wide performance limits than others, so signal receivers (e.g., potential mates and rivals) may be able to assess among-individual variation in song performance. Among song type variation in performance metrics indicates that some song types are more challenging to sing than others.

### Trade-offs and performance constraints

We found strong evidence of three acoustic trade-offs at the note level. The note-level scatterplots with sharply-defined, positively sloping boundaries comprise what we believe to be the most compelling acoustic evidence of performance constraints in bird song to date (Fig. 4, ESM-1-3). Individuals’ scatterplots and slope estimates showed that each male was subject to the trade-offs that constrain vocal production. Inter-individual variation in slope estimates for the note length vs. gap length and gap BW vs. gap length comparisons suggests the novel hypothesis that high quality individuals are subject to weaker trade-offs (lower slopes) than are low quality individuals.

The observed trade-off between note length and gap length supports the hypothesis that respiratory performance constrains song structure (Figs. 4a, d). Relative to short notes, long notes require birds to expire more air, necessitating longer subsequent mini-breaths to replenish the bird’s air supply in preparation for the next note. A similar pattern was previously identified in male serin (*Serinus serinus*) and dark-eyed junco (*Junco hyemalis*) songs (Mota and Cardoso 2001; Cardoso et al. 2007). The lower boundary for the note length vs. gap length distribution bulges downward at short note lengths in the pooled plot and in many of the individual plots (Figs. 4a, ESM-1). We hypothesize that this bulge represents notes that are short enough for birds to sing with pulsatile expiration (Hartley and Suthers 1989; Mota and Cardoso 2001). We do not yet know whether Adelaide’s warbler receivers respond to variation in respiratory performance, but there is evidence that other species do. Male skylarks (*Alauda arvensis*) increase the sound density of their songs when challenged with playback (Geberzahn and Aubin 2014b). Male dusky warblers (*Phylloscopus fuscatus*) that sing with consistently high amplitude live longer and enjoy extra-pair paternity benefits (Forstmeier et al. 2002). Similarly, male zebra finches (*Taeniopygia guttata*) that sing with higher sound density are preferred by females (Holveck and Riebel 2007).

We interpret the results from the note BW vs. note length and gap BW vs. gap length comparisons as evidence that frequency modulation speed is constrained (Figs. 4b, c, e, f). At the limit of performance, large frequency changes require more time than do small frequency changes. Podos (1997) first described a constraint on the speed of FM in general, and Geberzahn and Aubin (2014a) identified the constraint on unvoiced FM in skylarks. We believe that the present study is the first to specifically characterize a constraint on voiced FM. The apparent physiological basis of both constraints is that the magnitude of FM scales with the magnitude of physical change in the vocal apparatus and the vocal apparatus requires time to reconfigure itself (Suthers 2004). The speed of FM may be constrained by the bird’s ability to coordinate the various component of the vocal apparatus (brain, left and right syrinx, upper vocal tract, etc.; reviewed in Podos et al. 2009) that are involved in frequency modulation. Alternatively, a limitation on any component could limit the entire system. For example, bill size constrains singing speed in various taxa (Westneat et al. 1993; Podos 2001; Derryberry et al. 2012). There is widespread evidence that signal receivers attend to variation in FM speed (Drăgănoiu et al. 2002; Ballentine 2004; Illes et al. 2006; Cramer and Price 2007; DuBois et al. 2009; DuBois et al. 2011; Moseley et al. 2013; Vehrencamp et al. 2013; Phillips and Derryberry 2017a, 2017b).

We also found evidence of constrained performance at the song level. All four song-level quantile regressions were statistically significant, but the visual data from the length vs. TR and length vs. PoS comparisons were not compelling (Figs. 5c, d, e, f). We conclude that there is insufficient evidence to support the hypotheses that song length (or the combination of length and other variables) advertises singers’ motor abilities. Singing does not require much more oxygen than resting, and mini-breaths between notes may permit birds to escape respiratory constraints on song length (Oberweger and Goller 2001; Suthers and Zollinger 2004). Another study that found no evidence of constrained song length did find that birds sang longer songs when they were vocally interacting with other males (Cardoso et al. 2009). The authors concluded that ‘the length of songs is a plastic trait that varies with social context in a way that seems to reflect overall motivation or singing intensity’ (p. 904). A similar dynamic may be at play in our study system.

We found robust evidence that TR trades off against mean BW, replicating Podos’s (1997) canonical finding in sparrows (see Podos et al. 2016 for a general review), as well as a previous finding in other taxa (Wilson et al. 2014), including wood-warblers (Cardoso and Hu 2011). We originally believed that the TR vs. mean BW trade-off was caused by constraints on FM speed during voiced notes and unvoiced gaps. However, the modest correlations between TR vs. mean BW deviations and the mean deviations from note BW vs. note length (r = 0.223) and gap BW vs. gap length (r = 0.310) suggest that note-level FM constraints may not fully explain the TR vs. mean BW constraint. We conclude that TR vs. BW trade-offs probably emerge from some combination of constraints to voiced FM, unvoiced FM, and note repetition rate. The TR vs. PoS analysis provides further evidence that note repetition rate is constrained.

We tentatively conclude that the balance of evidence supports the existence of a TR vs. PoS trade-off, such that fast trills include more silence than do slow trills. The regression results were marginally significant (p = 0.0356), and visual analysis indicated negative trends in the pooled data, and some evidence of negative trends in the individual data (Figs. 5g, h, ESM-7). Although other studies have examined metrics similar to PoS (Forstmeier et al. 2002; Leadbeater et al. 2005; Holveck and Riebel 2007; Cardoso et al. 2009), we believe that this is the first study to show a relationship between PoS and TR. The physiological basis of this trade-off is probably similar to the respiratory constraint responsible for the note length vs. gap length trade-off. Indeed, the deviation scores from these two analyses are highly correlated (r = 0.886). To approach the upper boundary of the TR vs. PoS distribution, a bird must both minimize gap lengths, as in the note length vs. gap length trade-off, and also emit notes at a high rate.

At the limit of performance, the relationship between acoustic variables varied among individuals in both the TR vs. mean BW and TR vs. PoS comparisons. As in the note-level analysis, we propose that some individuals may be more robust to trade-offs than others, and that this variation may indicate quality.

### Comparing levels of analysis

The evidence for trade-offs at the note level was much stronger than the evidence at the song level. The boundaries of the note-level distributions were clearly defined, allowing us to see the shape of the boundary (e.g., the node in Fig. 4a). In contrast, the song-level boundaries were diffuse (compare Figs. 4 & 5). We believe that the main cause of this difference is that notes have a simpler structure than the songs they comprise. This simpler structure means that measured acoustic variables describe notes more completely than songs. Unmeasured variation among acoustic units contributes random error, making it harder to identify performance limits with song-level variables. Notes are also more numerous than songs, and large sample sizes permit more precise characterization of performance limits. It is often, but not always, possible to scale up from note-level performance to song-level performance (see the discussion of TR vs. PoS deviations above). We conclude that note-level analysis is a powerful approach for studying acoustic trade-offs, but it cannot completely replace song-level analyses.

### Methodological considerations

The present study revealed two important limitations to quantile regression. First, the line that it produces is not always parallel to the border of the point cloud, reducing our ability to accurately estimate performance limits and deviation scores. Part of the reason for the poor fit is that the quantile regression algorithm is influenced by points that are distant from the focal edge. This effect is most apparent in the gap BW vs. gap length scatterplot (Fig. 4c), in which the large number or long, low BW gaps influences the line to slope less steeply than the lower boundary. When the regression line does not parallel the edge of the distribution, deviation scores incorrectly weight their constituent variables. We overlooked this issue in the current study because our lines fit well enough to demonstrate trade-offs and approximate deviation scores, and because we did not want to introduce and justify a novel edge detection paradigm in a report that already contained many analyses. A second problem with the quantile regression analyses is that we found statistically significant quantile regressions even when plots did not appear to show strong evidence of performance limits (compare Figs. 5c-5f to the results of the corresponding quantile regressions). This problem was not unique to the *lqmm* package – standard quantile regressions were also statistically significant (unpublished analyses). Considering these limitations, we recommend that users supplement the results of quantile regression analyses with visual inspection of scatterplots and encourage the continued development of statistical methods for detecting and defining performance limits.

Our sample comprised many songs (n = 2879) and very many notes (n = 68,602), recorded from relatively few individuals (n = 9). This sampling scheme allowed us to characterize individuals’ distributions with high precision, especially at the note level. We were thus able to determine whether individuals were subject to trade-offs, and whether constraints varied among individuals. The large sample size of notes also resulted in dense point clouds that were amenable to visual assessment. Thus, we had a great deal of power to establish within individual patterns. Although the sample of individuals was modest, it was sufficient to test for population-level trade-offs without pseudoreplicating individuals because all individuals were subject to these note-level trade-offs.

### Repeatability of performance metrics

Most note-level deviation scores were repeatable by individual and by song type. ICC’s for individuals based on all songs indicated that 14-20% of variation in song performance can be attributed to individual differences. Whether receivers can differentiate among singers’ performance is an empirical question that is amenable to experimental investigation. Simulation models in which receivers sample varying numbers of songs from two singers and assess their relative performance, would also be useful. Estimates of repeatability by individual based on all songs tended to be considerably higher than those based on the averages for each song type within individual (Table 2). This difference has implications for both signallers and receivers.

Signallers do not sing all song types in their repertoires with equal frequency, but instead sing some song types more than others. Relative to a flat distribution of song types, the observed distribution of song types resulted in greater individual distinctiveness. This finding suggests the hypothesis that more skilled singers are able to sing demanding song types more often than are less skilled singers. Turning to receivers, our results indicate that estimating the average performance of all sampled songs would be a more efficient assessment strategy than would averaging for each song type and then averaging over song types. The former strategy is also less cognitively demanding because it includes fewer steps and requires less memory. Receivers’ assessment strategies require further investigation (Guilford and Dawkins 1991; Bateson and Healy 2005; Podos 2017).

Performance metrics were moderately to highly repeatable among song types (Tables 2 & 3). If deviation scores represent performance, then song types vary with respect to their performance demands. The deviation scores that measured FM speed (those that included BW and FEX) were highly repeatable by song type. That result was unsurprising because we used patterns of frequency modulation to classify song types (Fig. 2).

A previous study of male dark-eyed junco songs also found that repeatability estimates for song types were higher than repeatability estimates for individuals (Cardoso et al. 2009). That study averaged performance over song types within individuals before estimating individual ICC, and arrived at similar estimates to the comparable analysis in our study. Their estimates of song type repeatability were higher than ours, but that difference could be attributable to differences in song type scoring. The authors conclude that receivers could use acoustic performance to estimate singers’ quality, but go on to write, ‘the main conclusion from our results it that, because most of the variation in performance depends on the song type, a receiver that compares a few song types from different males is likely to obtain little information about performance differences between males’ (p. 905). In our study system, receivers would get considerably more information if they based their assessments on all songs (rather than averaging performance within song types and then within males). If high quality males sing more challenging song types, averaging performance within song type before calculating repeatability would underestimate receivers’ abilities to discern singers’ quality. Further, it is unlikely that Adelaide’s warbler receivers would only hear one or a few song types, because males rapidly cycle through their song types during dawn singing (Staicer 1991; Kaluthota et al. 2019).

### Correlations among performance metrics

Several pairs of performance metrics were positively correlated, suggesting that metrics are partially redundant or that different kinds of performance covary positively. It appears that note length vs. gap length deviation scores and gap BW vs. gap length deviation scores are highly correlated because short gaps generate low deviation scores in both comparisons. The correlation coefficient is inflated by the poor fit of the quantile regression line in the gap BW vs. gap length comparison (see above). Nevertheless, short gap lengths may indicate high quality with respect to both FM speed and respiratory performance. The high correlation between note BW vs. note length deviation and gap BW vs. gap length deviations may arise because short notes tend to be followed by short gaps, and vice-versa (as shown in the note length vs. gap length comparison).

At the level of the whole song, we found that FEX correlated strongly with various performance metrics that measure FM speed, but FEX was not strongly correlated with two metrics that seem to measure respiratory performance. Thus, FEX is useful for measuring overall FM speed, and note-level deviation scores are useful for parsing FM speed between voiced and unvoiced portions of the song, but FEX does not measure respiratory performance. The strong correlation between TR vs. PoS deviations and the deviation scores that involve gap length emphasizes the importance of short inter-note intervals for various performance metrics. Overall, we did not find evidence that FM performance trades off against respiratory performance.

### Skewness of performance metrics

Note-level deviation scores were positively skewed, as predicted by the hypothesis that the population has evolved under selection to sing near the physiological limits of performance (Table 2). We did not find this pattern with the song-level deviations (Table 3). The difference between patterns of skewness at the two levels of analysis may be attributable to the greater statistical noise in the song-level metrics. We tentatively conclude that the note-level skewness results support the performance constraint hypothesis and fail to support the constrained learning hypothesis, but equivalent data from other study systems are needed.

### Conclusions and future directions

Most research on performance constraints in bird song is based on acoustic trade-off analysis, but this paradigm has recently come under strong criticism (Kroodsma 2017b, 2017a). We responded to these criticisms with a study that included multiple comparisons at two levels, a note-level sample size that was nearly ten times larger than the largest comparable dataset (Geberzahn and Aubin 2014a), and novel statistical approaches to control for pseudoreplication and test for evidence of selection for high performance. We believe that this approach has produced the most compelling acoustic evidence yet that physiological limits constrain the structure of bird song.

We conclude that performance constraints on the speed of voiced FM, unvoiced FM, and respiration limit the acoustic structure of male Adelaide’s warbler songs. We hypothesize that this species sings trills of rapid frequency sweeps because of a history of sexual selection for FM and respiratory performance.

This study opens the door for future research on performance constraints in Adelaide’s warbler and other species. One critical question is whether male or female receivers attend to variation in one or more performance metrics in this population. It would be particularly interesting to know if they can accurately assess synthesized ‘singers’ with realistic performance ICC’s. Future studies should also examine the link between song performance and male phenotype, performance and among-individual variation in song type use, and how performance varies across contexts, including vocal interactions (Logue and Forstmeier 2008).

## Supporting information

Electronic Supplementary Material

## Acknowledgements

We would like to thank the people who helped us obtain and process recordings: Daniel Pereira and Josiris Rodriguez helped record birds; Paloma Sanchez-Jaureguí, Fabio L. Tarazona, Jorge Illanas, Krystal Medina, and Alfredo Lamela helped score recordings; Tony Shlakoff, Casandra Logue, Joshua Baez, Ruben Irizarry, and Alicia García processed song recordings in Luscinia; and Jesse McClure measured FEX. We also thank Dr. Robert Lachlan for his advice and technical support on Luscinia. Earlier versions of this manuscript were improved by comments from Hannes Schraft, Samantha Krause, Nolan Gooding, Todd Freeburg and two anonymous referees. This work was supported by a Discovery Grant from the Natural Sciences and Engineering Research Council of Canada (RGPIN-2015-06553) to D.M.L., and two NSERC undergraduate summer research assistantships to J.S. and B.W.

## Declaration

The authors have no conflicts of interest to declare.

## References

Ballentine B. 2004. Vocal performance influences female response to male bird song: an experimental test. Behavioral Ecology. 15(1):163–168.

Barr DJ, Levy R, Scheepers C, Tily HJ. 2013. Random effects structure for confirmatory hypothesis testing: Keep it maximal. Journal of Memory and Language. 68(3):255–278.

Bateson M, Healy SD. 2005. Comparative evaluation and its implications for mate choice. Trends in Ecology and Evolution. 20(12):659–664.

Bradbury J, Vehrencamp SL. 2011. Principles of Animal Communication: Second edition. Sunderland, Massachusetts: Sinauer.

Burnham KP, Anderson DR, Huyvaert KP. 2010. AIC model selection and multimodel inference in behavioral ecology: some background, observations, and comparisons. Behav Ecol Sociobiol. 65(1):23–35.

Byers J. 1997. American pronghorn: social adaptations and the ghosts of predators past. University of Chicago Press.

Byers J, Hebets E, Podos J. 2010. Female mate choice based upon male motor performance. Animal Behaviour. 79(4):771–778.

Cade BS, Noon BR. 2003. A gentle introduction to quantile regression for ecologists. Frontiers in Ecology and the Environment. 1(8):412–420.

Cardoso GC. 2013. Using frequency ratios to study vocal communication. Animal behaviour. 85(6):1529–1532.

Cardoso GC. 2017. Advancing the inference of performance in birdsong. Animal Behaviour. 125:e29–e32.

Cardoso GC, Atwell JW, Ketterson ED, Price TD. 2007. Inferring performance in the songs of dark-eyed juncos (*Junco hyemalis*). Behavioral Ecology. 18(6):1051–1057.

Cardoso GC, Atwell JW, Ketterson ED, Price TD. 2009. Song types, song performance, and the use of repertoires in dark-eyed juncos (*Junco hyemalis*). Behavioral Ecology. 20(4):901–907.

Cardoso GC, Hu Y. 2011. Birdsong performance and the evolution of simple (rather than elaborate) sexual signals. The American naturalist. 178(5):679–686.

Clark CJ. 2009. Courtship dives of Anna’s hummingbird offer insights into flight performance limits. Proceedings of the Royal Society B: Biological Sciences. 276(1670):3047–3052.

Cramer ERA, Price JJ. 2007. Red-winged blackbirds *Ageliaus phoeniceus* respond differently to song types with different performance levels. Journal of Avian Biology. 38(1):122–127.

Derryberry EP, Seddon N, Claramunt S, Tobias JA, Baker A, Aleixo A, Brumfield RT. 2012. Correlated evolution of beak morphology and song in the neotropical woodcreeper radiation. Evolution: International Journal of Organic Evolution. 66(9):2784–2797.

Drăgănoiu TI, Nagle L, Kreutzer M. 2002. Directional female preference for an exaggerated male trait in canary (*Serinus canaria*) song. Proceedings of the Royal Society of London B: Biological Sciences. 269(1509):2525–2531.

DuBois AL, Nowicki S, Searcy WA. 2009. Swamp sparrows modulate vocal performance in an aggressive context. Biology letters. 5(2):163–165.

DuBois AL, Nowicki S, Searcy WA. 2011. Discrimination of vocal performance by male swamp sparrows. Behav Ecol Sociobiol. 65(4):717–726.

Forstmeier W, Kempenaers B, Meyer A, Leisler B. 2002. A novel song parameter correlates with extra-pair paternity and reflects male longevity. Proceedings of the Royal Society of London B: Biological Sciences. 269(1499):1479–1485.

Geberzahn N, Aubin T. 2014a. Assessing vocal performance in complex birdsong: A novel approach. BMC Biology. 12:1–9.

Geberzahn N, Aubin T. 2014b. How a songbird with a continuous singing style modulates its song when territorially challenged. Behav Ecol Sociobiol. 68(1):1–12.

Geraci M. 2014. Linear quantile mixed models: the lqmm package for Laplace quantile regression. Journal of Statistical Software. 57(13):1–29.

Guilford T, Dawkins MS. 1991. Receiver psychology and the evolution of animal signals. Animal Behaviour. 42(1):1–14.

Hartley RS, Suthers RA. 1989. Airflow and pressure during canary song: direct evidence for mini-breaths. Journal of Comparative Physiology A. 165(1):15–26.

Hedley RW, Logue DM, Benedict L, Mennill DJ. 2018. Assessing the similarity of song-type transitions among birds: evidence for interspecies variation. Animal Behaviour. 140:161–170.

Hoese WJ, Podos J, Boetticher NC, Nowicki S. 2000. Vocal tract function in birdsong production: experimental manipulation of beak movements. Journal of Experimental Biology. 203(12):1845–1855.

Holveck M-J, Riebel K. 2007. Preferred songs predict preferred males: consistency and repeatability of zebra finch females across three test contexts. Animal Behaviour. 74(2):297–309.

Illes AE, Hall ML, Vehrencamp SL. 2006. Vocal performance influences male receiver response in the banded wren. Proceedings of the Royal Society of London B: Biological Sciences. 273(1596):1907–1912.

Kaluthota CD, Medina OJ, Logue DM. 2019. Quantifying song categories in Adelaide’s Warbler (*Setophaga adelaidae*). Journal of Ornithology.1–11.

Kroodsma D. 2017a. Birdsong ‘performance’studies: A sad commentary. Animal Behaviour. 133:209–210.

Kroodsma D. 2017b. Birdsong performance studies: a contrary view. Animal Behaviour. 125:e1–e16.

mLachlan R. 2007. Luscinia: a bioacoustics analysis computer program.

Larimer JL, Dudley R. 1995. Accelerational implications of hummingbird display dives. The Auk. 112(4):1064–1066.

Leadbeater E, Goller F, Riebel K. 2005. Unusual phonation, covarying song characteristics and song preferences in female zebra finches. Animal Behaviour. 70(4):909–919.

Logue DM, Forstmeier W. 2008. Constrained performance in a communication network: implications for the function of song-type matching and for the evolution of multiple ornaments. The American naturalist. 172(1):34–41.

Moseley DL, Lahti DC, Podos J. 2013. Responses to song playback vary with the vocal performance of both signal senders and receivers. Proceedings of the Royal Society B: Biological Sciences. 280(1768):20131401.

Mota PG, Cardoso GC. 2001. Song organisation and patterns of variation in the serin (*Serinus serinus*). Acta Ethologica. 3(2):141–150.

Murai M, Backwell PR. 2006. A conspicuous courtship signal in the fiddler crab *Uca perplexa*: female choice based on display structure. Behav Ecol Sociobiol. 60(5):736–741.

Nowicki S, Westneat M, Hoese W. 1992. Birdsong: motor function and the evolution of communication. Seminars in the Neurosciences. 4(6):385–390.

Oberweger K, Goller F. 2001. The metabolic cost of birdsong production. Journal of Experimental Biology. 204(19):3379–3388.

Phillips JN, Derryberry EP. 2017a. Equivalent effects of bandwidth and trill rate: support for a performance constraint as a competitive signal. Animal behaviour. 132:209–215.

Phillips JN, Derryberry EP. 2017b. Vocal performance is a salient signal for male–male competition in White-crowned Sparrows. The Auk: Ornithological Advances. 134(3):564–574.

Plummer EM, Goller F. 2008. Singing with reduced air sac volume causes uniform decrease in airflow and sound amplitude in the zebra finch. Journal of Experimental Biology. 211(1):66–78.

Podos J. 1997. A performance constraint on the evolution of trilled vocalizations in a songbird family (Passeriformes: Emberizidae). Evolution; international journal of organic evolution. 51(2):537–551.

Podos J. 2001. Correlated evolution of morphology and vocal signal structure in Darwin’s finches. Nature. 409:185–188.

Podos J. 2017. Birdsong performance studies: reports of their death have been greatly exaggerated. Animal Behaviour. 125:e17–e24.

Podos J, Lahti DC, Moseley DL. 2009. Vocal performance and sensorimotor learning in songbirds. Advances in the Study of Behavior. 40:159–195.

Podos J, Moseley DL, Goodwin SE, McClure J, Taft BN, Strauss AV, Rega-Brodsky C, Lahti DC. 2016. A fine-scale, broadly applicable index of vocal performance: frequency excursion. Animal behaviour. 116:203–212.

Podos J, Nowicki S. 2004. Beaks, adaptation, and vocal evolution in Darwin’s finches. AIBS Bulletin. 54(6):501–510.

mRudis B, Bolker B, Schulz J, Kothari A, Sidi J. 2017. ggalt: extra coordinate systems,‘Geoms’, statistical transformations, scales and fonts for ‘ggplot2’.

Schraft HA, Medina OJ, McClure J, Pereira DA, Logue DM. 2017. Within-day improvement in a behavioural display: wild birds ‘warm up’. Animal Behaviour. 124:167–174.

Smith JM, Harper D. 2003. Animal signals. Oxford University Press.

Staicer C. 1991. The role of male song in the socioecology of the tropical resident Adelaide’s warbler (Dendroica adelaidae). University of Massachusetts.

Stoffel MA, Nakagawa S, Schielzeth H. 2017. rptR: Repeatability estimation and variance decomposition by generalized linear mixed-effects models. Methods in Ecology and Evolution. 8(11):1639–1644.

Suthers RA. 2004. How birds sing and why it matters. Nature’s music: the science of birdsong Elsevier Academic Press, San Diego.272–295.

Suthers RA, Zollinger SA. 2004. Producing song: the vocal apparatus. Annals of the New York Academy of Sciences. 1016(1):109–129.

Team R. 2015. RStudio: Integrated Development for R.

Toms JD. 2010. Adelaide’s Warbler (Setophaga adelaidae). version 1.0. Ithaca, New York, USA: Cornell Lab of Ornithology; [accessed 2018 July 4].

Vehrencamp SL, Yantachka J, Hall ML, de Kort SR. 2013. Trill performance components vary with age, season, and motivation in the banded wren. Behav Ecol Sociobiol. 67(3):409–419.

Westneat MW, Long J, Hoese W, Nowicki S. 1993. Kinematics of birdsong: functional correlation of cranial movements and acoustic features in sparrows. Journal of Experimental Biology. 182(1):147–171.

Wickham H, Chang W. 2008. ggplot2: An implementation of the Grammar of Graphics. R package version 07, URL: http://CRANR-projectorg/package=ggplot2.

Wilson DR, Bitton PP, Podos J, Mennill DJ. 2014. Uneven sampling and the analysis of vocal performance constraints. The American naturalist. 183(2):214–228.

