## Supplementary material for "An analysis of avian vocal performance at the note and song levels": Electronic Supplementary Material

**Table ESM-1:** Random intercept and slope estimates for individuals from quantile regression analyses (tau = 0.1) on acoustic properties of Adelaide's warbler notes.

| x | y | individual | intercept | slope |
|---|---|---|---|---|
| note length | gap length | DDLb | 8.64 | 0.32 |
| | | KYK | 10.84 | 0.30 |
| | | LgRLg | 9.96 | 0.24 |
| | | LgWV | 9.64 | 0.28 |
| | | OWO | 12.82 | 0.14 |
| | | PDP | 11.36 | 0.23 |
| | | RbRbO | 9.28 | 0.27 |
| | | RDY | 8.76 | 0.24 |
| | | ROLb | 11.99 | 0.18 |
| note BW | note length | DDLb | -1.39 | 13.51 |
| | | KYK | -0.29 | 10.41 |
| | | LgRLg | 11.42 | 6.71 |
| | | LgWV | 0.90 | 13.81 |
| | | OWO | 12.03 | 4.53 |
| | | PDP | 19.51 | 4.04 |
| | | RbRbO | 8.30 | 9.96 |
| | | RDY | -7.61 | 14.48 |
| | | ROLb | 16.24 | 5.95 |
| gap BW | gap length | DDLb | 17.88 | 2.17 |
| | | KYK | 16.72 | 2.38 |
| | | LgRLg | 16.68 | 1.31 |
| | | LgWV | 13.99 | 4.26 |
| | | OWO | 23.07 | -2.93 |
| | | PDP | 18.87 | 0.98 |
| | | RbRbO | 18.48 | 1.21 |

| | | | |
|---|---|---|---|
| | RDY | 13.90 | 1.79 |
| | ROLb | 16.74 | 1.20 |

**Table ESM-2.** Results of quantile regression analyses (tau = 0.9) on acoustic properties of Adelaide's warbler songs.

| x | y | individual | intercept | slope |
|---|---|---|---|---|
| trill rate | mean BW | DDLb | 3.34 | -0.09 |
| | | KYK | 2.74 | -0.03 |
| | | LgRLg | 2.63 | -0.02 |
| | | LgWV | 4.04 | -0.16 |
| | | OWO | 3.57 | -0.09 |
| | | PDP | 2.68 | -0.03 |
| | | RbRbO | 3.13 | -0.07 |
| | | RDY | 3.21 | -0.07 |
| | | ROLb | 3.41 | -0.08 |
| song length | trill rate | DDLb | 14.69 | -0.0007 |
| | | KYK | 14.89 | -0.0004 |
| | | LgRLg | 15.09 | -0.0006 |
| | | LgWV | 14.84 | -0.0007 |
| | | OWO | 15.70 | -0.0005 |
| | | PDP | 14.89 | -0.0008 |
| | | RbRbO | 15.36 | -0.0011 |
| | | RDY | 14.38 | 0.0001 |
| | | ROLb | 14.97 | -0.0008 |
| song length | PoS | DDLb | 69.09 | -0.0026 |
| | | KYK | 69.92 | -0.0041 |
| | | LgRLg | 69.58 | -0.0020 |
| | | LgWV | 69.97 | -0.0025 |
| | | OWO | 67.81 | -0.0014 |

| | | | | |
|---|---|---|---|---|
| | | PDP | 70.50 | -0.0023 |
| | | RbRbO | 69.38 | -0.0016 |
| | | RDY | 71.05 | -0.0028 |
| | | ROLb | 70.44 | -0.0018 |
| trill rate | PoS | DDLb | 73.56 | -0.89 |
| | | KYK | 73.90 | -1.00 |
| | | LgRLg | 74.33 | -0.81 |
| | | LgWV | 74.27 | -0.90 |
| | | OWO | 75.24 | -0.86 |
| | | PDP | 74.30 | -0.79 |
| | | RbRbO | 74.09 | -0.75 |
| | | RDY | 74.95 | -0.84 |
| | | ROLb | 74.38 | -0.70 |

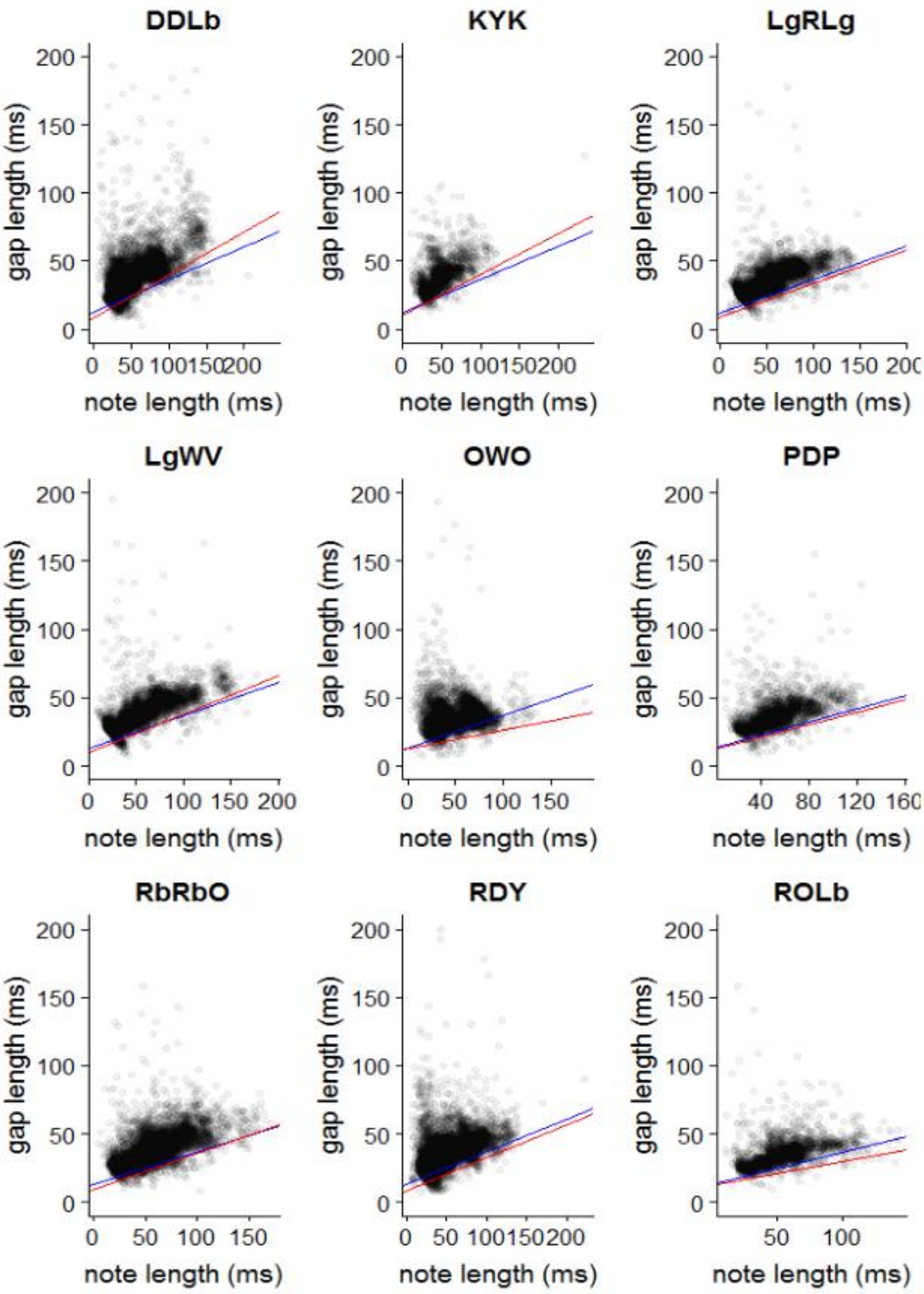


**Figure ESM-1.** Scatterplots of note length vs. gap length for nine male Adelaide's warblers. Blue lines represent the tenth quantile regression line for the pooled data. Red lines represent the tenth quantile regression for the focal individual. Titles indicate the birds' IDs.

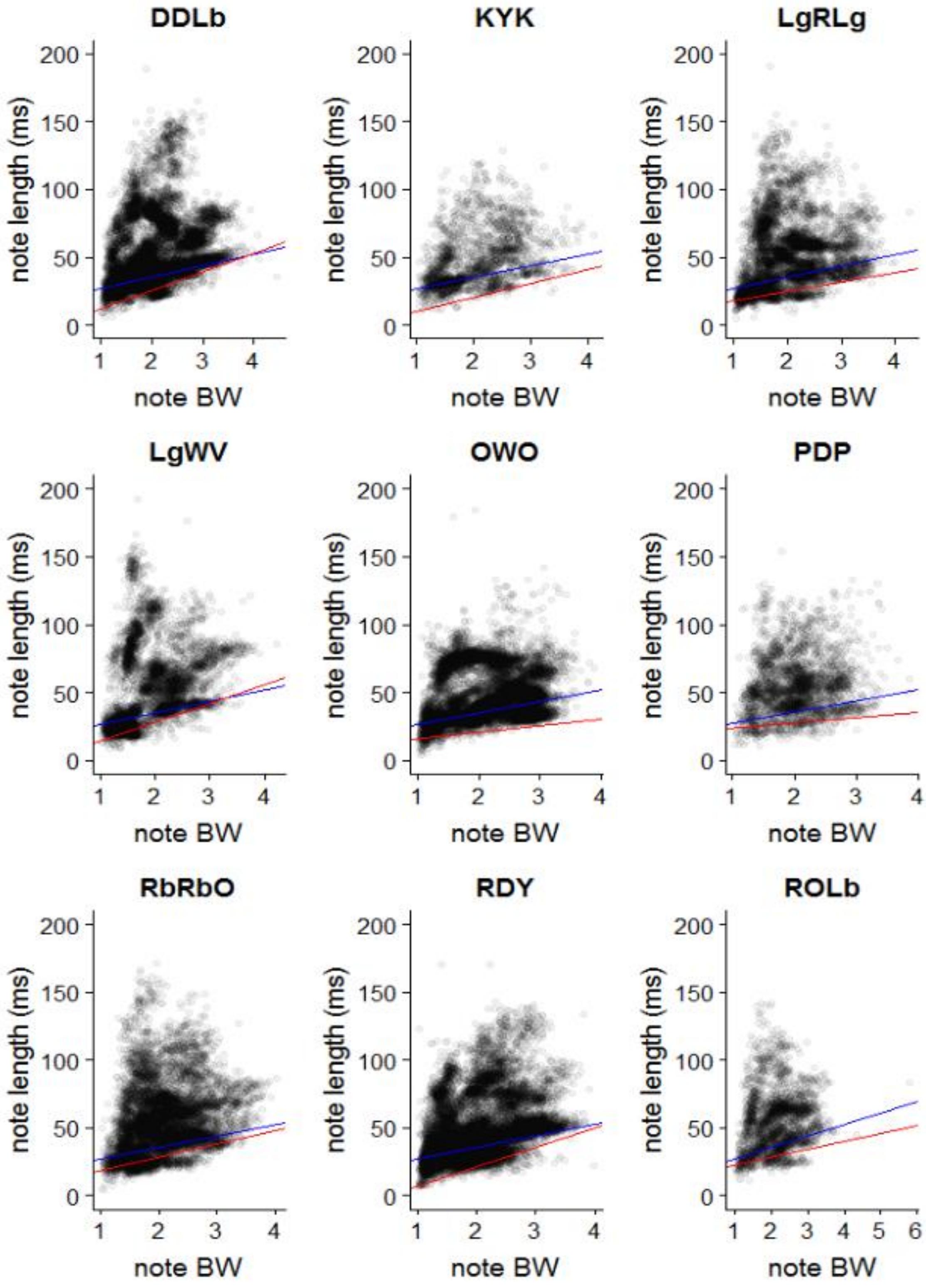


**Figure ESM-2.** Scatterplots of note bandwidth vs. gap length for nine male Adelaide's warblers. Blue lines represent the tenth quantile regression line for the pooled data. Red lines represent the tenth quantile regression for the focal individual. Titles indicate the birds' IDs.

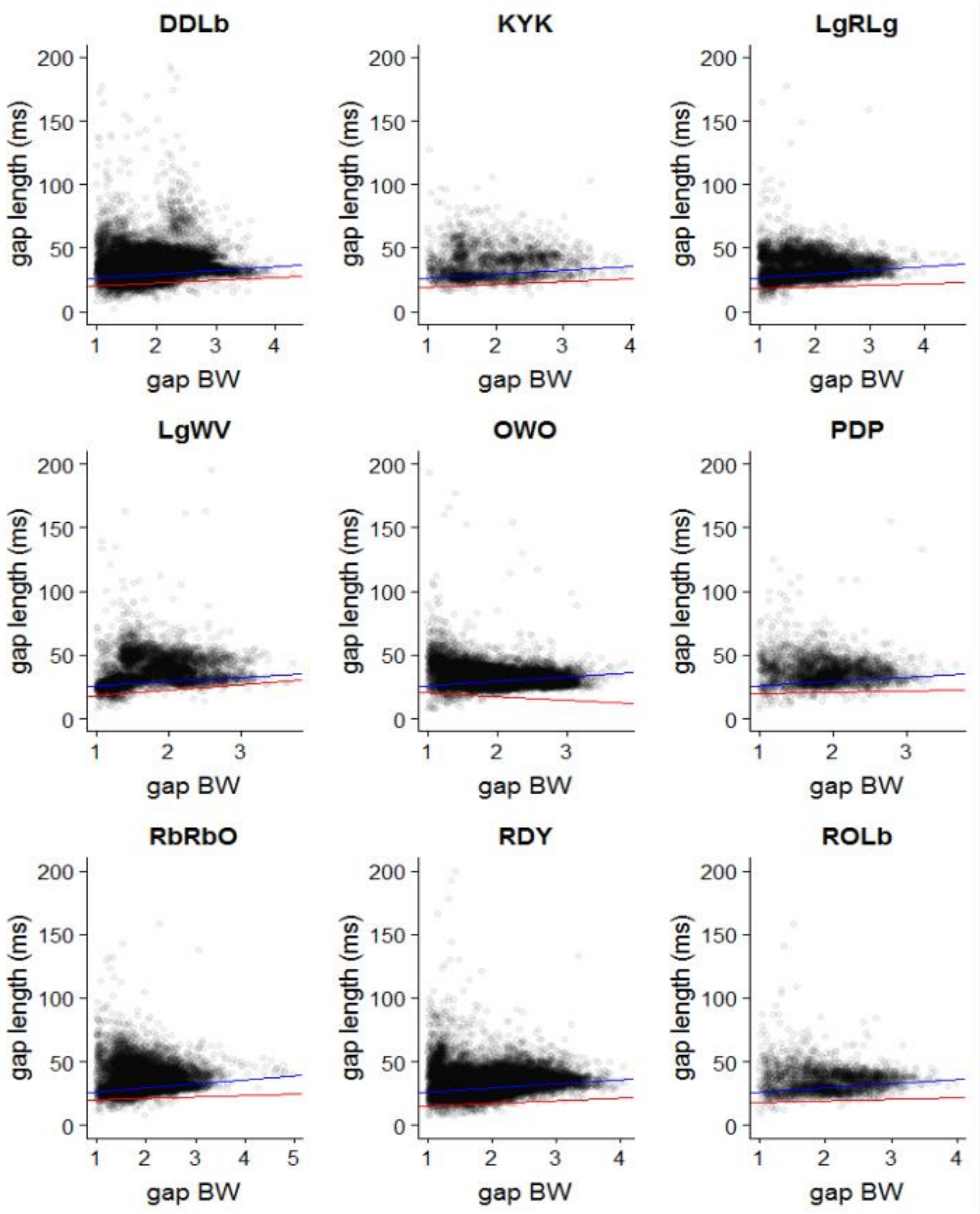


**Figure ESM-3.** Scatterplots of gap bandwidth vs. gap length for nine male Adelaide's warblers. Blue lines represent the tenth quantile regression line for the pooled data. Red lines represent the tenth quantile regression for the focal individual. Titles indicate the birds' IDs.

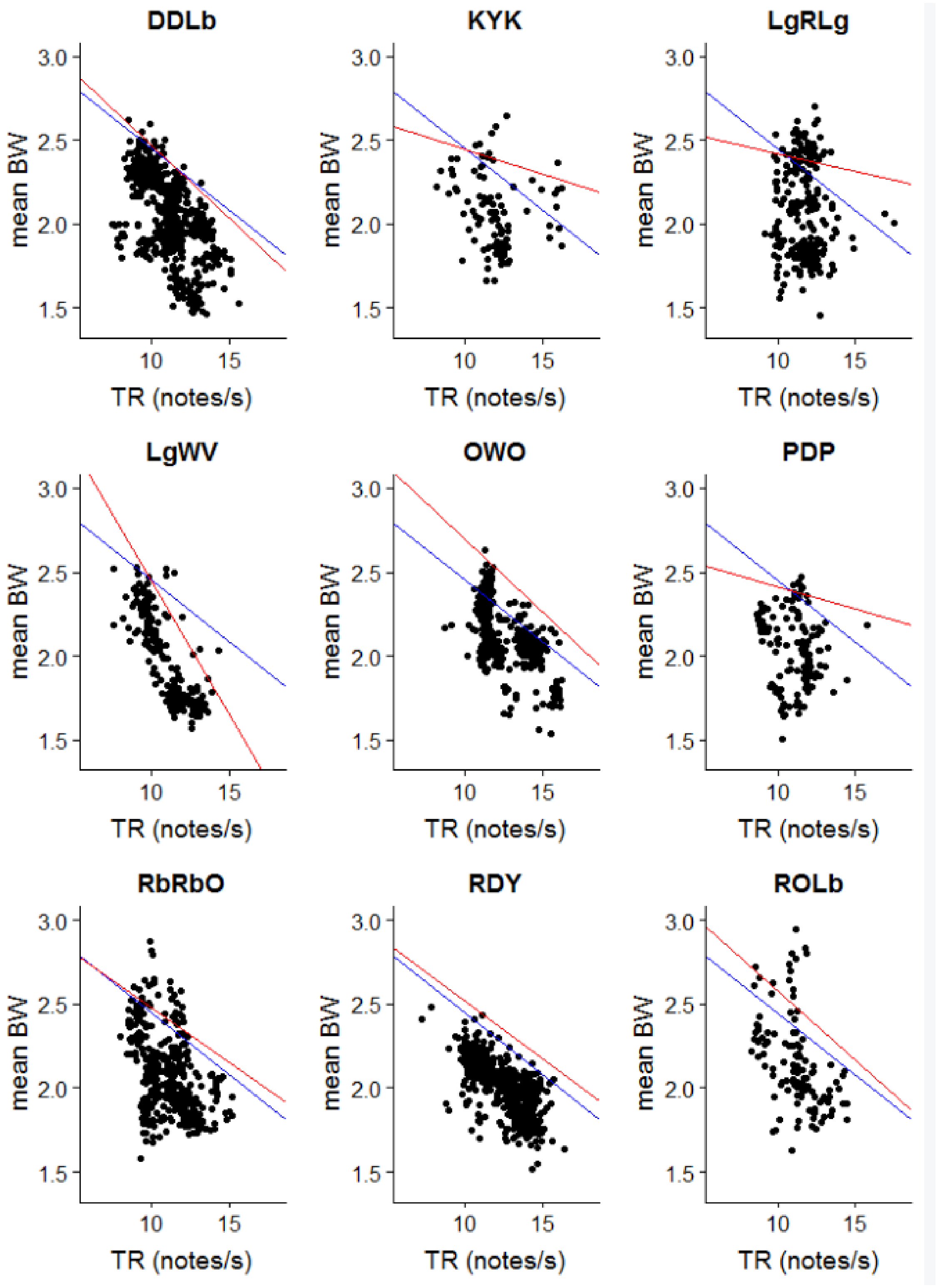


**Figure ESM-4.** Scatterplots of trill rate vs. mean bandwidth for nine male Adelaide's warblers. Blue lines represent the tenth quantile regression line for the pooled data. Red lines represent the tenth quantile regression for the focal individual. Titles indicate the birds' IDs.

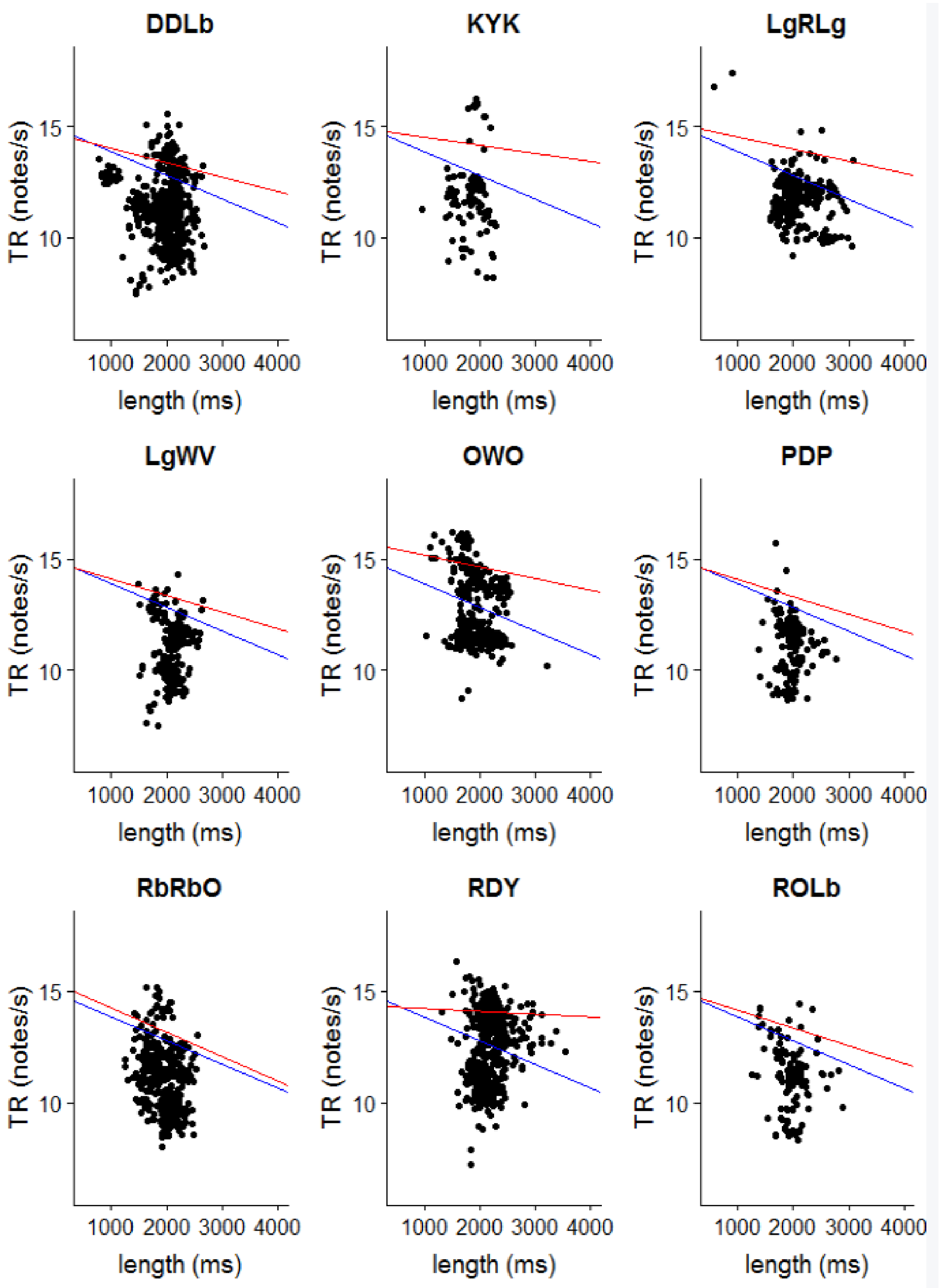


**Figure ESM-5.** Scatterplots of song length vs. trill rate for nine male Adelaide's warblers. Blue lines represent the tenth quantile regression line for the pooled data. Red lines represent the tenth quantile regression for the focal individual. Titles indicate the birds' IDs.

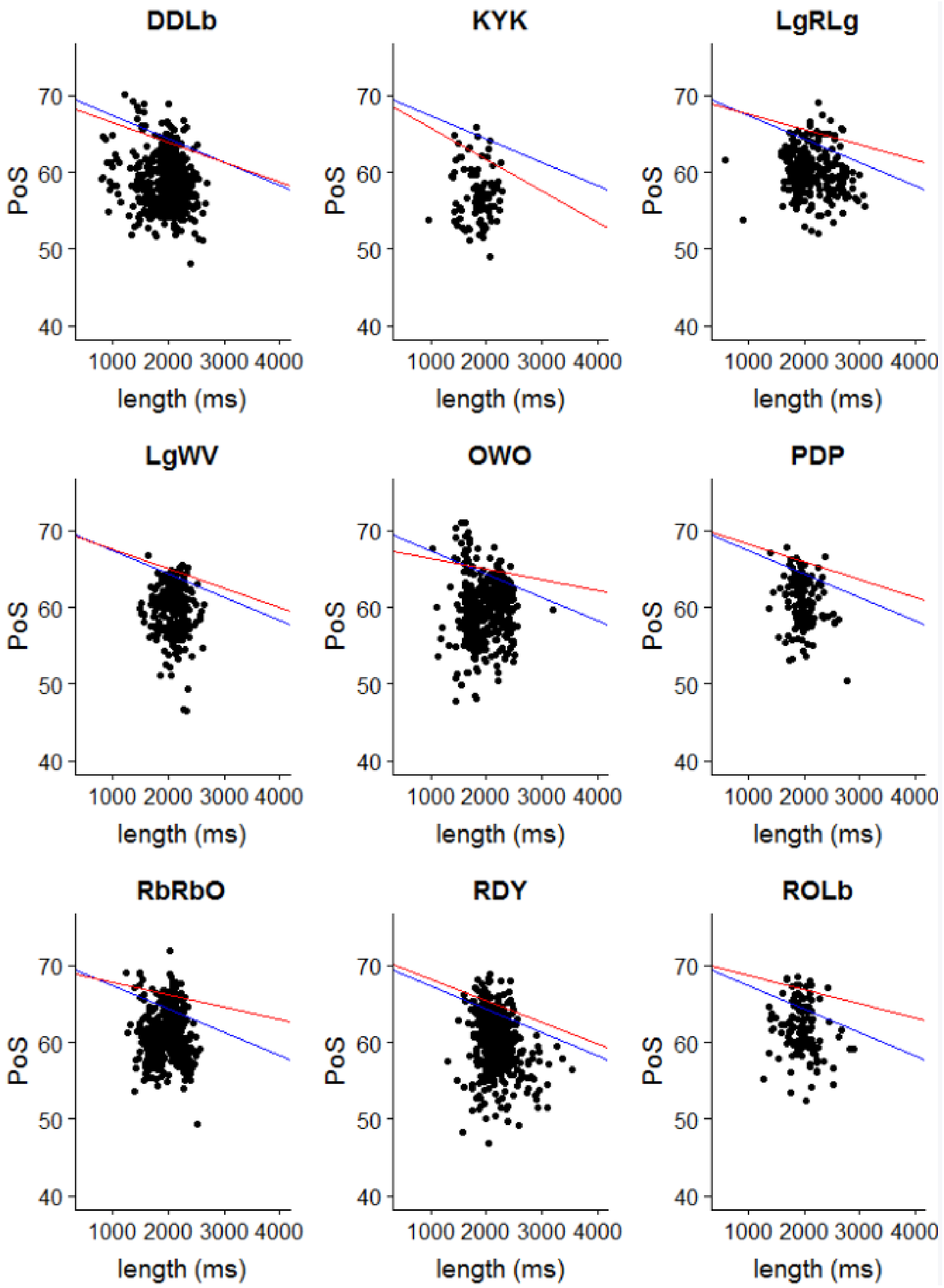


**Figure ESM-6.** Scatterplots of song length vs. percent of sound for nine male Adelaide's warblers. Blue lines represent the tenth quantile regression line for the pooled data. Red lines represent the tenth quantile regression for the focal individual. Titles indicate the birds' IDs.

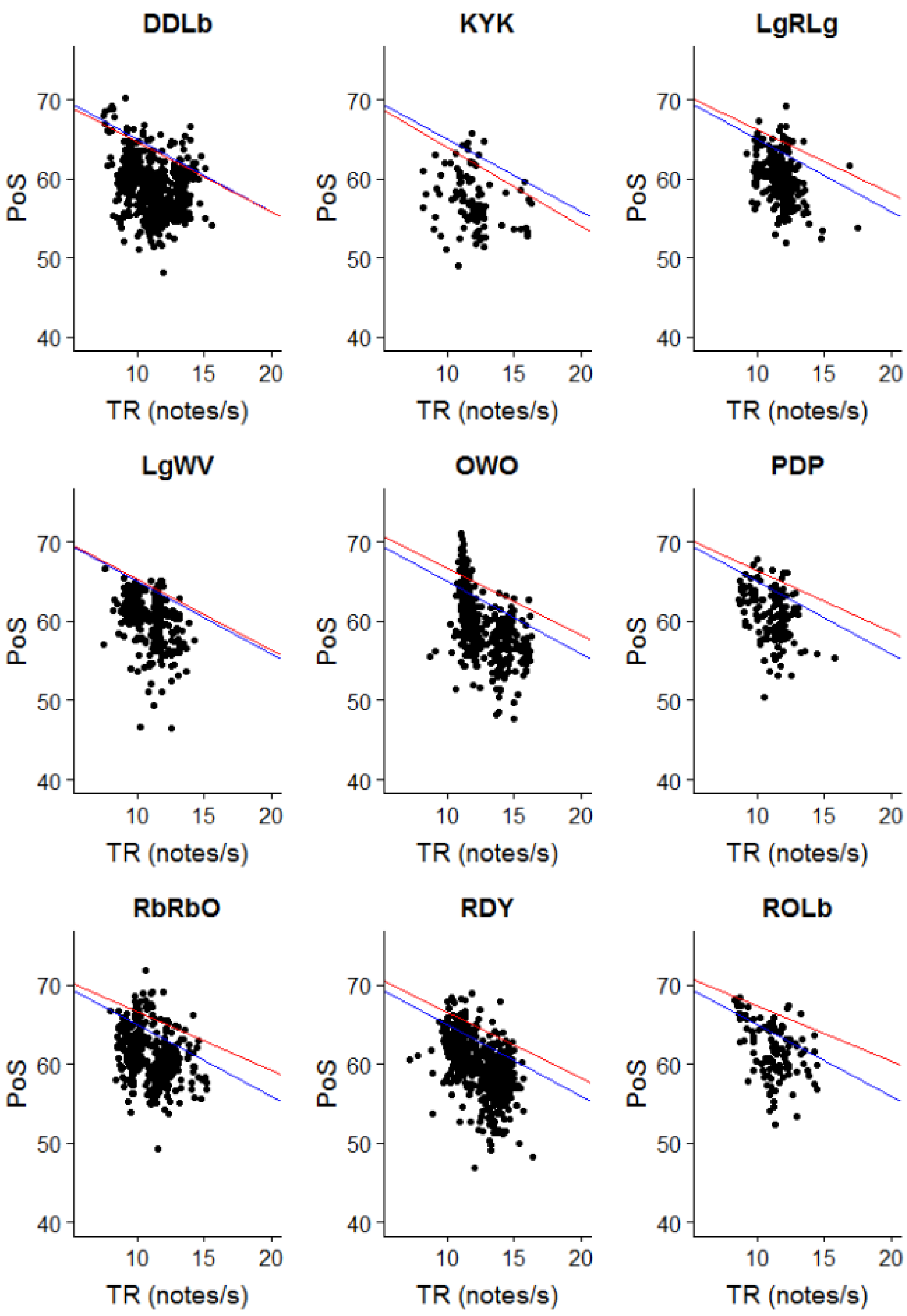


**Figure ESM-7.** Scatterplots of trill rate vs. percent of sound for nine male Adelaide's warblers. Blue lines represent the tenth quantile regression line for the pooled data. Red lines represent the tenth quantile regression for the focal individual. Titles indicate the birds' IDs.
